# The Effect of COVID-19 on the Postdoctoral Experience: a comparison of pre-pandemic and pandemic surveys

**DOI:** 10.1101/2021.11.19.468693

**Authors:** Andréanne. Morin, Britney A. Helling, Seetha. Krishnan, Laurie E. Risner, Nykia D. Walker, Nancy B. Schwartz

## Abstract

In the interest of advocating for the postdoctoral community in the United States, we present results from survey data collected before and during the COVID-19 pandemic on the same population of postdocs. In 2019, 5,929 postdocs in the US completed a comprehensive survey, and in 2020, a subset completed a follow-up survey several months into the pandemic. The results show that the pandemic has substantially impacted postdocs’ mental health and wellness irrespective of gender, race, citizenship, or other identities. Postdocs also reported a significant impact on their career trajectories and progression, reduced confidence in achieving career goals, and negative perceptions of the job market compared to pre-COVID-19. International postdocs also reported experiencing distinct stressors due to the changes in immigration policy. Notably, having access to Postdoctoral Associations and Postdoctoral Offices positively impacted postdocs’ overall well-being and helped mitigate the personal and professional stresses and career uncertainties caused by the pandemic.

**Graphical Abstract:** Graphical Abstract of survey responses to: Why or how has your research been disrupted or not disrupted due to the pandemic? Overall, postdocs responded with feelings of loss of control as the pandemic was acting upon them and taking away their ability to complete their work.

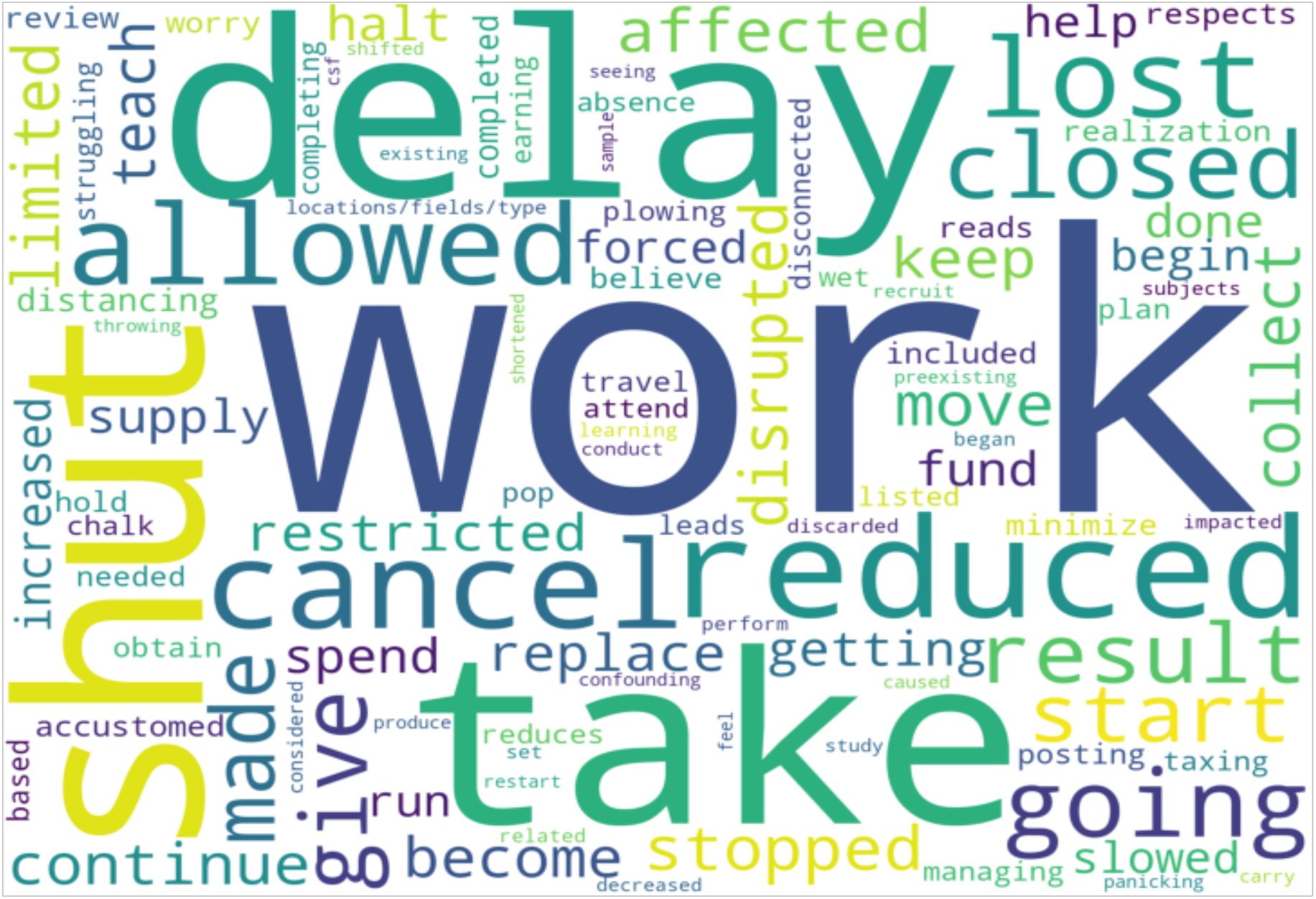

## Introduction

Often unknown to those outside of the scientific community and overlooked by their own institutions compared to faculty and students, postdocs have long been referred to as the invisible component of the University^1^. Typically, postdocs lack job security as they are funded by research grants tied to an individual faculty member and are subjected to annual contracts; they receive lower pay in comparison to non-academic peers in government or industry; and frequently lack employee-type benefits such as paid family leave^2,3^. The COVID-19 pandemic has made these situations worse for postdocs^4–7^.

The impact of the pandemic on postdocs is not unlike the severe and far-reaching effects the COVID-19 pandemic has had worldwide. In the US alone, significant job loss, educational disparities, and elevated mental health issues have dramatically affected the workforce^7–9^; such that the detrimental impact on the global economy may extend through the next decade^10^. There has been a similar adverse effect on the biomedical workforce^11–14^. Although the financial impact of COVID-19 on scientific productivity has not yet been fully realized, the NIH estimates a $16 billion loss because of delayed research^15^. In fact, the Bureau of Labor Statistics reported the largest decline in college and university employment since the 1950s^5,16,17^. Furthermore, numerous universities retracted or deferred new faculty job offers, leaving postdocs, who are the source of future academics, to either consider different career paths or extend their current postdoc positions^18^.

One report in Nature has addressed the impact of the pandemic on the STEM postdoc population^19^. This report indicates that nearly two-thirds of postdocs surveyed believed that their long-term career prospects were negatively affected by the COVID-19 pandemic; roughly 8 out of 10 postdocs reported that the pandemic had hampered their ability to conduct experiments and collect data, and more than half had difficulty communicating with supervisors and colleagues.

We have long been interested in the postdoctoral experience in the US with respect to career choices, mentorship, grantsmanship, and gender disparities and in 2016 released the first comprehensive survey of postdocs^20^ since 2005^21^. Our extensive study of over 7,500 postdocs from 351 institutions assessed the factors that influenced postdoc satisfaction and career plans^20^. We conducted a second survey from mid to late 2019 to continue tracking these aspects of the postdoc experience over time. This updated survey queried >6,000 postdocs from various institutions nationwide. As the effects of the COVID-19 pandemic began to be felt widely, a follow-up survey was conducted in the Fall of 2020 on a subset (n=1,942) of the 2019 survey respondents to assess the impact of the pandemic on the postdoc trainee population.

Here we present a comparison of survey data collected before and during the COVID-19 pandemic on the same group of postdocs working in the US. We investigated the impact of the pandemic on mental health and wellness, changes in their career trajectories and progression, and their confidence in achieving their career goals. Due to government policy changes enacted during the pandemic that affected international travel, immigration, and visa access, we also looked at specific challenges that the pandemic had on international postdocs working in the US. Finally, we investigated the impact of COVID-19 on the availability of wellness and mental health resources, as well as the role that institutional Postdoctoral Associations and Postdoctoral Offices had on postdocs’ overall well-being during the pandemic, as these critical factors have not been previously explored.

## Results

In 2019 (June to December), we conducted a national survey to assess the postdoctoral experience in the US. The goal of this initial survey was to serve as an update and to expand upon our national survey conducted in 2016^20^. In the early months of 2020, the consequences of the COVID-19 pandemic began to impact daily life across the United States. To understand the effects of the pandemic in the context of the postdoctoral experience, we re-surveyed a subset of postdocs who completed the 2019 survey between October 1 and November 3 of 2020. This follow-up survey allowed us to query the same population before and during the pandemic to assess its consequences more directly.

### Demographics

In 2019, 6,292 respondents participated in our national postdoc survey, of which 5,929 identified as postdocs in the US. These respondents were 58% female, 41% male and 0.4% non-binary/third gender (**Figure 1A**). Regarding race and ethnicity, 60% of the respondents were white, 27% were Asian, and 13% were from underrepresented minority backgrounds (URMs; because some racial and ethnic groups were small, we combined individuals into these three main categories for analyses - see Methods for a full description and **Supplementary file 3** for a more granular description) (**Figure 1B**). US citizens or Permanent Residents (PR; referred to as US citizens/PR throughout this manuscript) made up 53% of the respondents, and 47% were international postdocs working in the US on temporary visas (J1, H1B, TN, F1, F1-OTP, E3 visas) (**Figure 1C**). The majority (55%) of postdocs were 30-34 years old (**Figure 1D**), and most respondents were in their first (39%) or second (29%) year of their postdoctoral training (**Figure 1E**). Respondents were from various disciplines, mostly within life sciences (48%), followed by medicine, physical sciences, engineering, psychology, environmental sciences, and social sciences, among other research areas. (**Figure 1F**).

**Figure 1:**
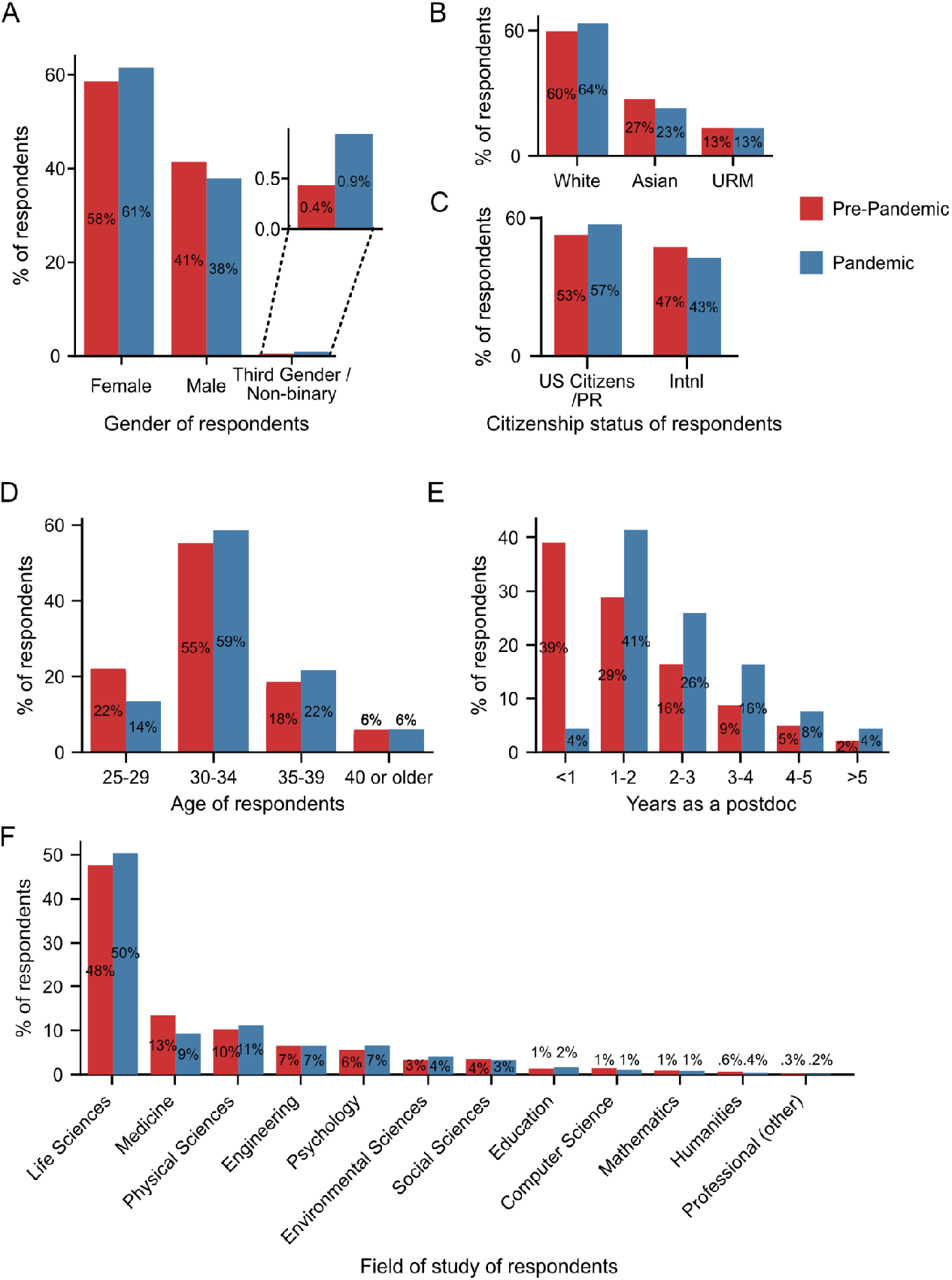
Pre-pandemic and pandemic survey demographics. **A**. More self-identified female and third gender/non-binary and fewer self-identified male respondents completed the pandemic survey (n=1,698) compared to the pre-pandemic survey (n=5,805; Chi-squared test, p=0.0023, χ2 = 12.2). **B**. The majority of respondents were white in both the pre-pandemic (n=5,649) and pandemic surveys (n=1,673), with an increase in white and a decrease in Asian respondents in the pandemic survey compared to the pre-pandemic survey (Chi-squared test, p=0.0024, χ2 =12.1). **C**. The proportion of US citizens/PR respondents increased (Chi-squared test, p=0.0015, χ2 = 10.1; n pre-pandemic=5,813; n pandemic=1,702). **D-E**. As expected, the age of respondents (Chi-squared test, p=3.6×10^−14^, χ2 = 65.7; n pre-pandemic=5,825; n pandemic=1,714) **(D)** and the years of postdoc experience (Chi-squared test, p=4.3×10^−161^, χ2 = 755.8; n pre-pandemic=5,853; n pandemic=1,715) **(E)** both increased as we conducted the pandemic survey with a subset of the pre-pandemic respondents almost one year after the initial survey. **F**. The majority of respondents were in the life sciences with a statistically significant decrease in responses from those in the field of medicine in the pandemic survey (n=1,712) compared to the pre-pandemic survey (n=5,922; Chi-squared test, p=0.0012, χ2 = 32.47). PR: Permanent resident. Additional demographic information from the two surveys is shown in **Figure 1–figure supplement 1**

In October of 2020, 1,942 of the 6,292 respondents who participated in the 2019 survey, completed a follow-up survey assessing the effects of the COVID-19 pandemic. Of these, 1,722 (89%) were still in a postdoctoral position at a US institution. From here on, we refer to the 2019 survey as the pre-pandemic survey and the 2020 survey as the pandemic survey. Furthermore, in our analyses of current postdocs, we removed the 11% of respondents in the pandemic survey who were no longer in postdoctoral positions, however, we analyzed their career outcomes separately in **Figure 5**.

As shown in **Figure 1**, the demographics of the respondents to the pandemic survey largely mirrored those of the pre-pandemic survey. The number of respondents in the pandemic and pre-pandemic survey by US states is shown in **Figure 1–figure supplement 1A-B**. There were slightly more responses from individuals who identified as female (61% vs. 58%) and non-binary/third gender (0.9% vs. 0.4%), and fewer self-identified males (38% vs. 42%) in the pandemic survey compared to the pre-pandemic survey (**Figure 1A**). Race and ethnicity varied between the pre-pandemic and pandemic survey respondents, with a 4% increase in the proportion of respondents who identify as white and a corresponding 4% decrease in the respondents who identify as Asian. No differences were observed between the proportion of URMs (13%; **Figure 1B; Figure 1– figure supplement 1C**) or in identity groups (i.e., disability, LGBTQ, and veterans) (**Figure 1–figure supplement 1D**). When analyzed by citizenship, there was an increase in respondents who were US citizens/PR (53% pre-pandemic vs. 57% pandemic) and a corresponding decrease in international respondents (47% pre-pandemic vs. 43% pandemic) (**Figure 1C**). Given that we conducted the pandemic survey within a sub-population of those in the pre-pandemic survey at a later date, the age of the pandemic respondents was higher than the pre-pandemic respondents, and as expected, they were more advanced in their postdoc tenure (**Figure 1D-E**). There was a significant decrease in respondents in the field of medicine (13% pre-pandemic and 9% pandemic), while there was no significant change in the representation of any other field (**Figure 1F**). Lastly, there was a significant increase in access to a PDO (65.6% pre-pandemic vs 70% pandemic), which was mainly due to an increased awareness, but no differences in term of access to a PDA (**Figure 1–figure supplement 1F-G**).

### COVID-19 Impact

To directly assess the effects of COVID-19 on postdocs, we queried three general areas: stressors during the pandemic, institutional response to the pandemic, and ability to meet basic needs. In an open-ended question enquiring about the main stressors during the pandemic, postdocs indicated that their main stressors were a combination of work, family, and emotional burdens, as shown by the word cloud analysis of the responses (**Figure 2A-B**). Individual responses showed how postdocs experienced different types of burdens. Parents and caregivers faced the burden of “being a full time [*sic*] postdoc and staying home with two kids” or caring for a loved one who was/is struggling with COVID-19. As one postdoc indicated, “my girlfriend has been recovering from COVID-19 since March. It’s a grueling process to watch and support.” A large number of postdocs also indicated that work progress was more difficult due to “getting research done within limited shifts and hours” and an overall fear of “loss of productivity”. Many international postdocs were concerned about their visas and one respondent even indicated that the international office at their institution told them “…you will lose your job if you leave the country for any reason and are not a resident.” **Table 1** includes additional representative responses.

**Figure 2:**
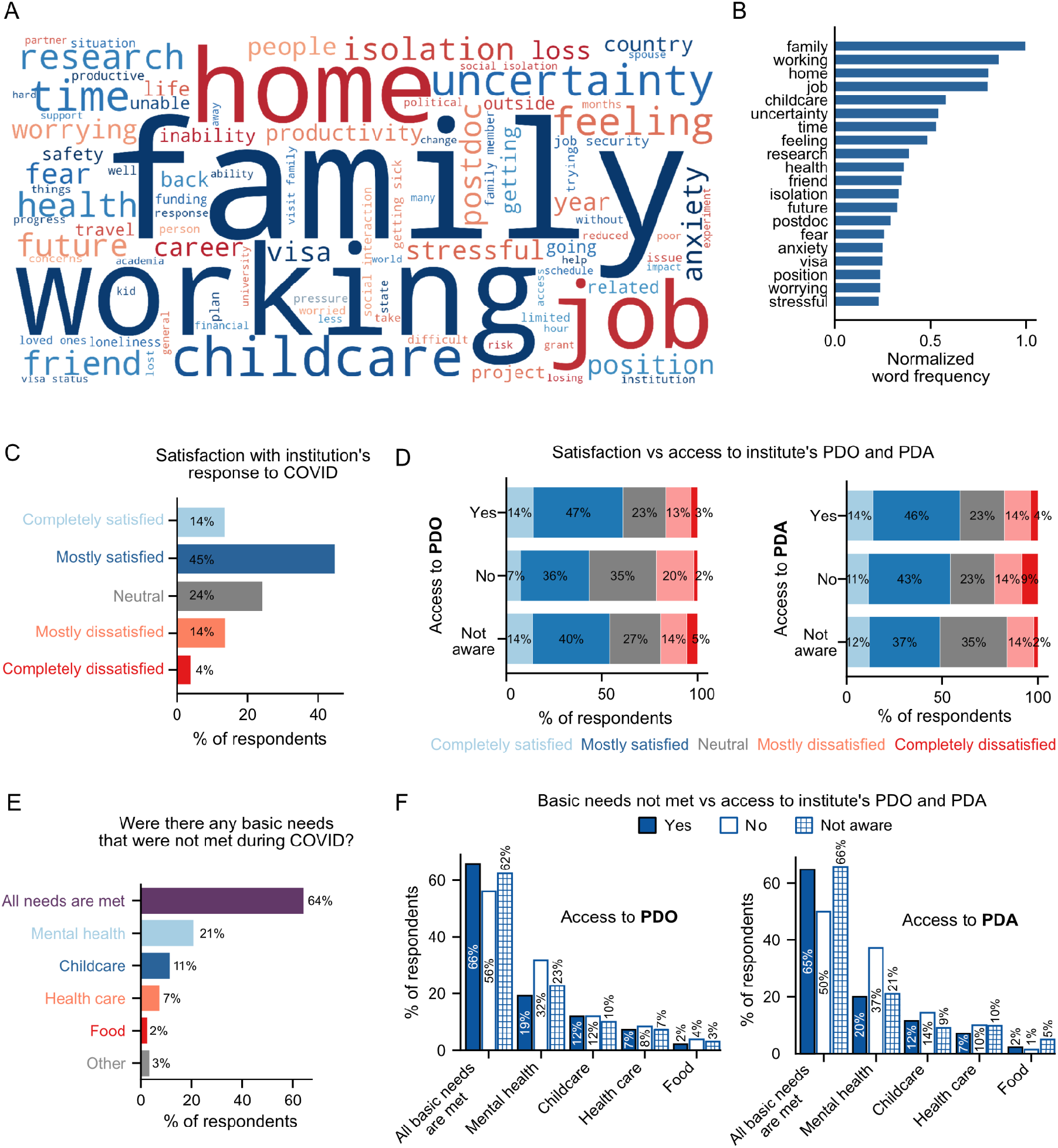
Impact of the pandemic on postdocs and effect of institutional support. **A-B**. Word cloud of postdocs’ main stressors during the COVID-19 pandemic **(A)** and distribution of the most frequently used words **(B). C**. Satisfaction with the institution’s response to COVID-19 (n=1,718). **D**. Satisfaction with the institution’s response to COVID-19 was higher in postdocs that had access to a PDO compared to the ones that did not (ordinal logistic regression OR=1.75 [95% CI; 1.23-2.48], p=0.0018) or those unaware whether their institution had a PDO (ordinal logistic regression OR=1.24 [95% CI; 1.01-1.53], p=0.044; n=1,700). No significant differences were observed by access to PDA (n=1,707). **E**. Basic needs that were not met during the pandemic (n=1,676). See **Figure 2–figure supplement 1** for breakdown by race/ethnicity groups. **F**. Having access to a PDO significantly impacted having mental health needs met (Chi-squared test, p=0.005, χ2 = 10.6, n=1,660). Having access to a PDA significantly impacted having all their basic needs (Chi-squared test, p=0.039, χ2 = 6.5) or meeting their mental health needs (Chi-squared test, p=0.0026, χ2 = 11.9; n=1,665).

**Table 1.**
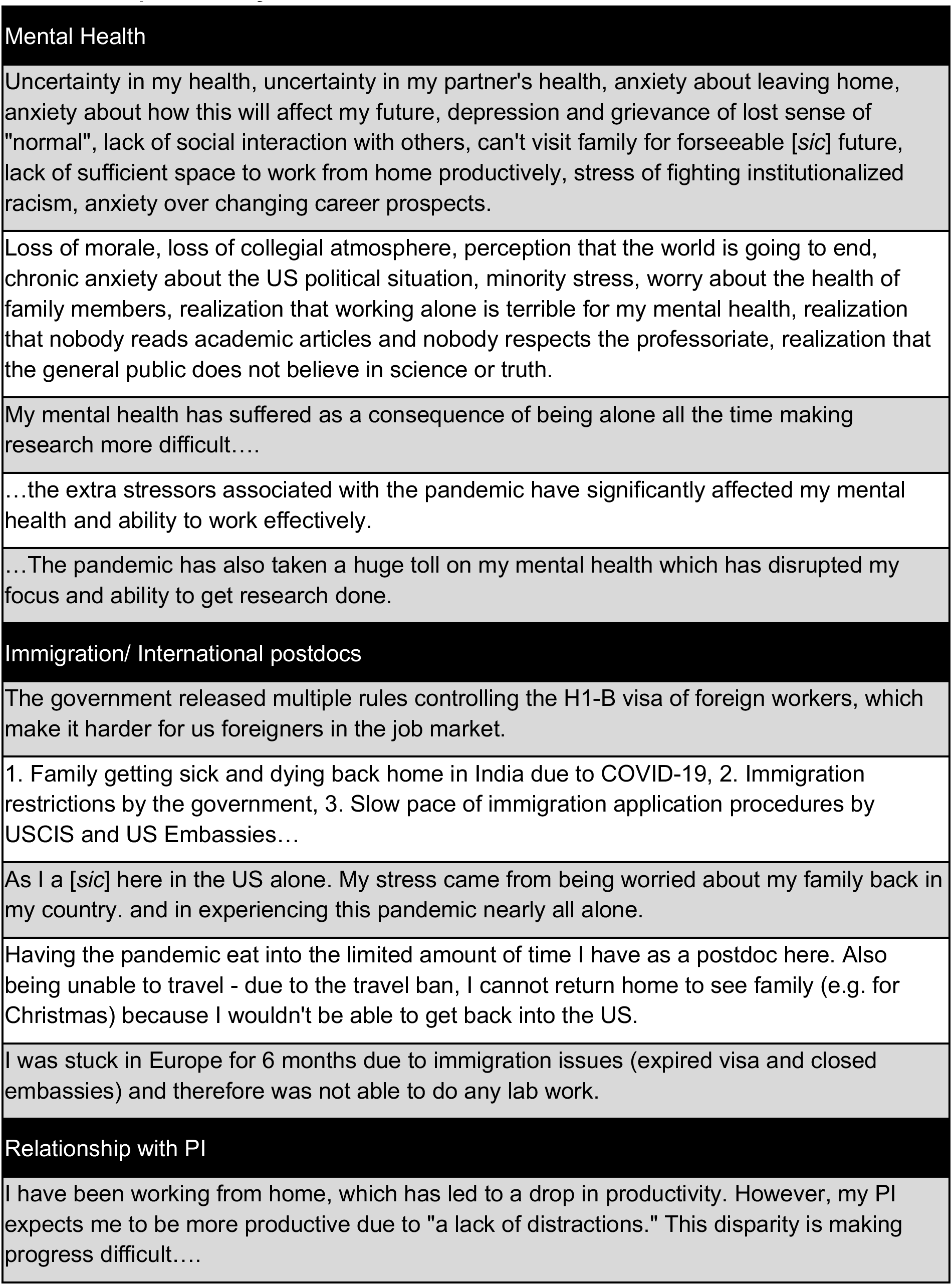
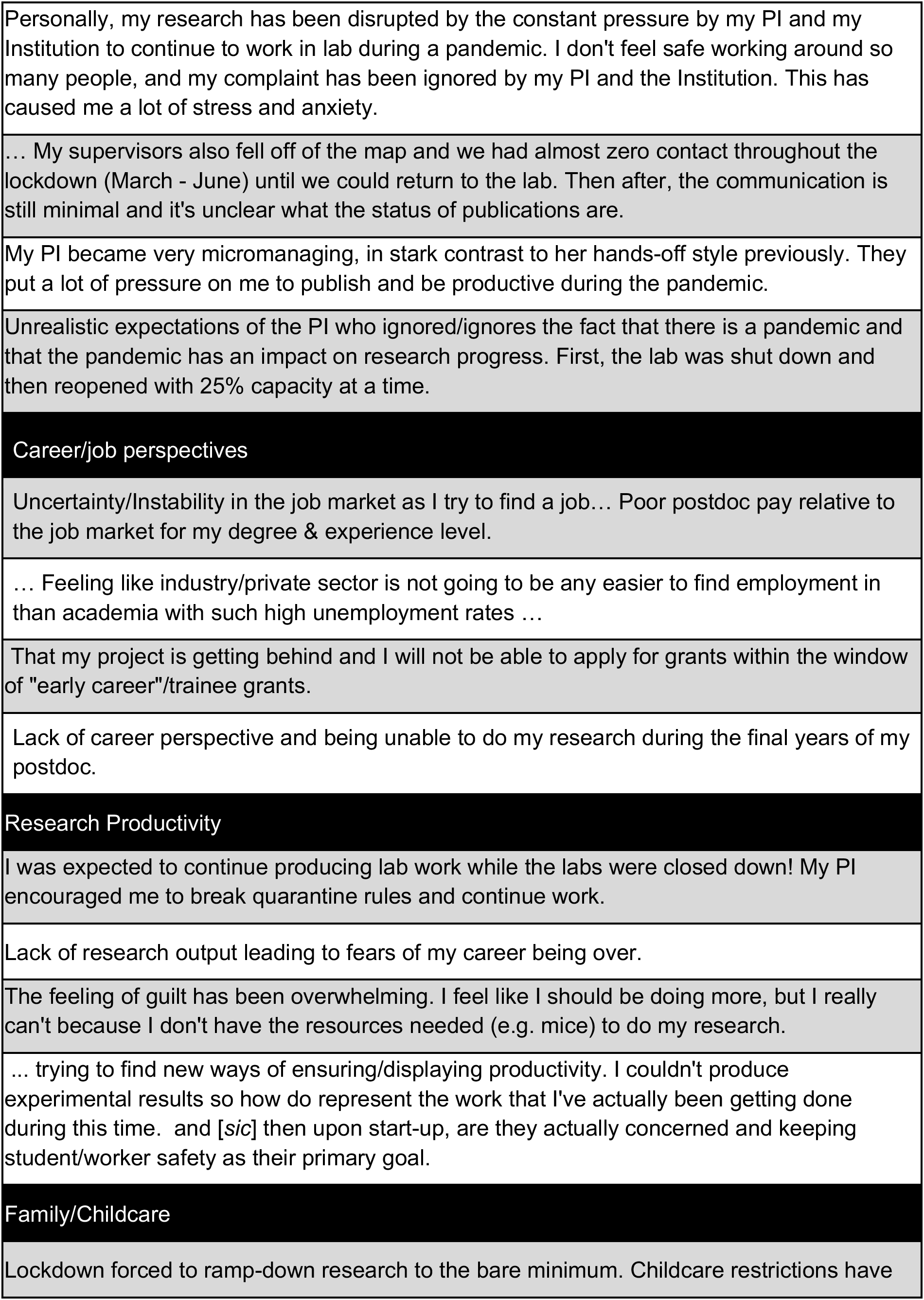
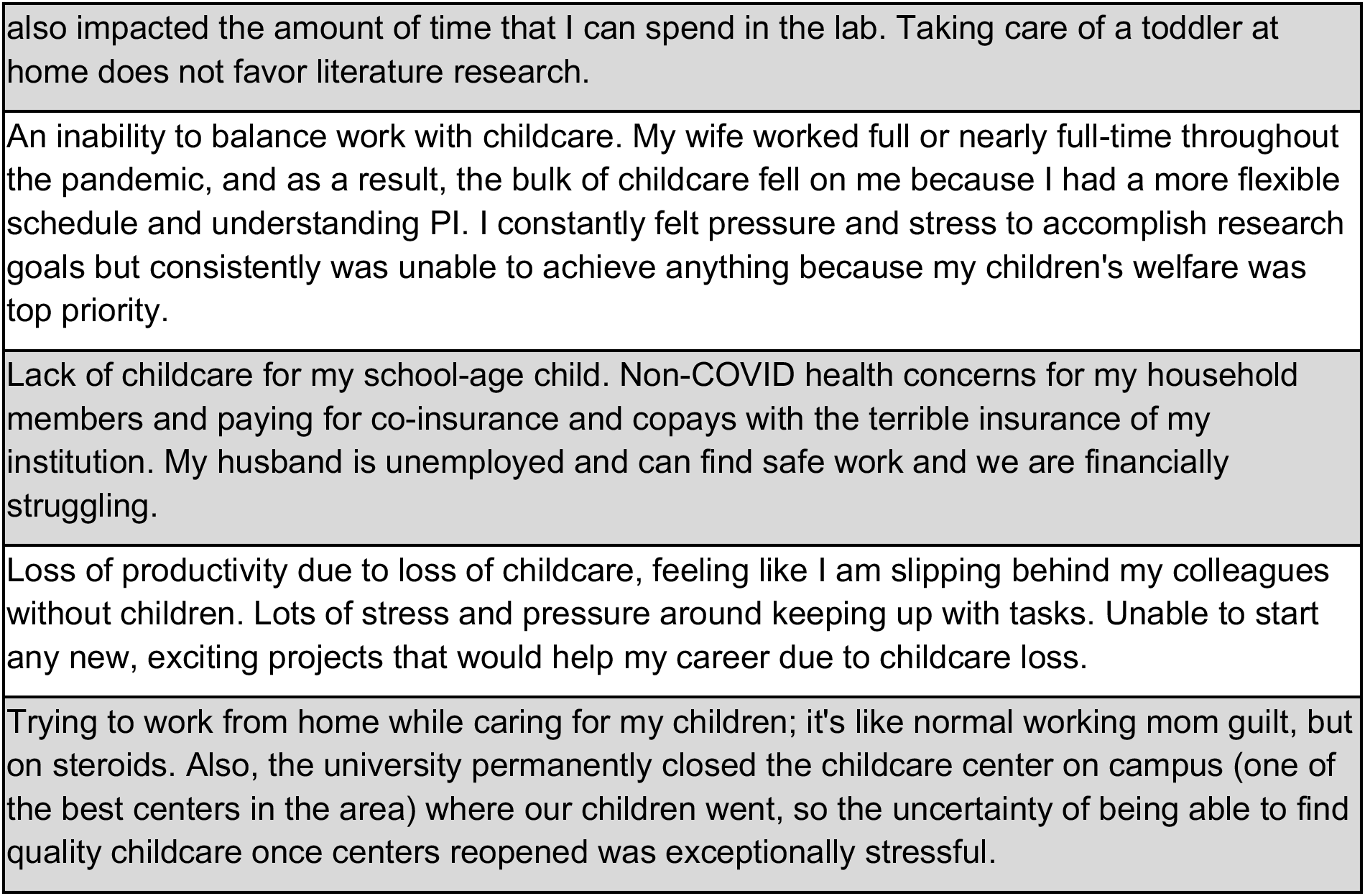
Responses to open-ended questions on pandemic-related stresses and impact on research productivity.

| Mental Health |  |
| --- | --- |
|  | Uncertainty in my health, uncertainty in my partner's health, anxiety about leaving home, anxiety about how this will affect my future, depression and grievance of lost sense of "normal", lack of social interaction with others, can't visit family for foreseeable [sic] future, lack of sufficient space to work from home productively, stress of fighting institutionalized racism, anxiety over changing career prospects. |
|  | Loss of morale, loss of collegial atmosphere, perception that the world is going to end, chronic anxiety about the US political situation, minority stress, worry about the health of family members, realization that working alone is terrible for my mental health, realization that nobody reads academic articles and nobody respects the professoriate, realization that the general public does not believe in science or truth. |
|  | My mental health has suffered as a consequence of being alone all the time making research more difficult.... |
|  | ...the extra stressors associated with the pandemic have significantly affected my mental health and ability to work effectively. |
|  | ...The pandemic has also taken a huge toll on my mental health which has disrupted my focus and ability to get research done. |
| Immigration/ International postdocs |  |
|  | The government released multiple rules controlling the H1-B visa of foreign workers, which make it harder for us foreigners in the job market. |
|  | 1. Family getting sick and dying back home in India due to COVID-19, 2. Immigration restrictions by the government, 3. Slow pace of immigration application procedures by USCIS and US Embassies... |
|  | As I a [sic] here in the US alone. My stress came from being worried about my family back in my country. and in experiencing this pandemic nearly all alone. |
|  | Having the pandemic eat into the limited amount of time I have as a postdoc here. Also being unable to travel - due to the travel ban, I cannot return home to see family (e.g. for Christmas) because I wouldn't be able to get back into the US. |
|  | I was stuck in Europe for 6 months due to immigration issues (expired visa and closed embassies) and therefore was not able to do any lab work. |
| Relationship with PI |  |
|  | I have been working from home, which has led to a drop in productivity. However, my PI expects me to be more productive due to "a lack of distractions." This disparity is making progress difficult.... |

|  |
| --- |
| Personally, my research has been disrupted by the constant pressure by my PI and my Institution to continue to work in lab during a pandemic. I don't feel safe working around so many people, and my complaint has been ignored by my PI and the Institution. This has caused me a lot of stress and anxiety. |
| ... My supervisors also fell off of the map and we had almost zero contact throughout the lockdown (March - June) until we could return to the lab. Then after, the communication is still minimal and it's unclear what the status of publications are. |
| My PI became very micromanaging, in stark contrast to her hands-off style previously. They put a lot of pressure on me to publish and be productive during the pandemic. |
| Unrealistic expectations of the PI who ignored/ignores the fact that there is a pandemic and that the pandemic has an impact on research progress. First, the lab was shut down and then reopened with 25% capacity at a time. |
| **Career/job perspectives** |
| Uncertainty/Instability in the job market as I try to find a job... Poor postdoc pay relative to the job market for my degree & experience level. |
| ... Feeling like industry/private sector is not going to be any easier to find employment in than academia with such high unemployment rates ... |
| That my project is getting behind and I will not be able to apply for grants within the window of "early career"/trainee grants. |
| Lack of career perspective and being unable to do my research during the final years of my postdoc. |
| **Research Productivity** |
| I was expected to continue producing lab work while the labs were closed down! My PI encouraged me to break quarantine rules and continue work. |
| Lack of research output leading to fears of my career being over. |
| The feeling of guilt has been overwhelming. I feel like I should be doing more, but I really can't because I don't have the resources needed (e.g. mice) to do my research. |
| ... trying to find new ways of ensuring/displaying productivity. I couldn't produce experimental results so how do represent the work that I've actually been getting done during this time. and [sic] then upon start-up, are they actually concerned and keeping student/worker safety as their primary goal. |
| **Family/Childcare** |
| Lockdown forced to ramp-down research to the bare minimum. Childcare restrictions have |
| also impacted the amount of time that I can spend in the lab. Taking care of a toddler at home does not favor literature research. |
| An inability to balance work with childcare. My wife worked full or nearly full-time throughout the pandemic, and as a result, the bulk of childcare fell on me because I had a more flexible schedule and understanding PI. I constantly felt pressure and stress to accomplish research goals but consistently was unable to achieve anything because my children's welfare was top priority. |
| Lack of childcare for my school-age child. Non-COVID health concerns for my household members and paying for co-insurance and copays with the terrible insurance of my institution. My husband is unemployed and can find safe work and we are financially struggling. |
| Loss of productivity due to loss of childcare, feeling like I am slipping behind my colleagues without children. Lots of stress and pressure around keeping up with tasks. Unable to start any new, exciting projects that would help my career due to childcare loss. |
| Trying to work from home while caring for my children; it's like normal working mom guilt, but on steroids. Also, the university permanently closed the childcare center on campus (one of the best centers in the area) where our children went, so the uncertainty of being able to find quality childcare once centers reopened was exceptionally stressful. |

Next, we looked at the institutional response to COVID-19, which ranged from completely satisfied to completely unsatisfied. Most postdocs indicated that they were completely or mostly satisfied with their institution’s response to COVID-19 (59%) (**Figure 2C**). In particular, postdocs with access to a Postdoctoral Affairs Office (PDO; i.e., institutional entities, staffed by professionals and funded by institutions, that advocate for postdocs and promote initiatives to support postdoctoral training and professional development, and establish policies for compensation, benefits, term limits, eligibility, etc.) were significantly more satisfied than those who did not or were unaware of this institutional asset (**Figure 2D**). Moreover, there were no differences in satisfaction to their institution’s response between those with or without access to a Postdoctoral Affairs Association (PDA; i.e., institutional organizations composed of and managed by postdoctoral scholars that actively engage and represent the postdocs) (**Figure 2D**), or with respect to gender, citizenship status, race and ethnicity, or identity (ordinal logistic regression, p>0.05, data not shown). Notably, there was also a non-negligible portion (4%) of postdocs who indicated they were completely unsatisfied with their institution’s response to COVID-19, with one respondent commenting, “… my institution did almost NOTHING to ensure that faculty and staff can be safely back at work”.

Although the majority of postdocs indicated that all of their basic needs were met during the pandemic (64%), a significant portion (36%) indicated that their needs concerning mental health (21%), childcare (11%), healthcare (7%) and/or food (2%) were unmet (**Figure 2E**). Additionally, 3% of postdocs wrote in responses mentioning other unmet needs, including the inability to pay bills, exercise, loss of access to transportation, work safety, human connections, or loss of salary, retirement benefits, or annual raise. Furthermore, although the majority of postdocs indicated that all of their basic needs were met, the comments indicated that the pandemic had made meeting those needs more difficult: “My husband lost his job, and while we are not in danger of basic needs not being met it does change some things and adds additional stress”. Postdocs who had all of their basic needs met were more likely to have access to a PDA (65% (yes (access to a PDA)) and 50% (no (no access to a PDA)); **Figure 2F**). Furthermore, postdocs with access to a PDO or a PDA were less likely to have their mental health needs unmet (PDO: 32% (no) vs. 19% (yes); PDA: 37% (no) vs. 20% (yes), no differences were observed between those not aware and aware of a PDA or PDO at their institution, **Figure 2F**). Lastly, postdocs who identified as Asian (the majority of whom were international (76%)) were more likely than white postdocs to report unmet needs with respect to health care (12% vs. 5%) or food (5% vs. 1%) (**Figure 2–figure supplement 1A**). No differences were observed according to gender, identity, or URM status (Chi-squared test, p>0.05, data not shown).

Postdoc parents were particularly affected by pandemic-related shutdowns. While we did not directly inquire of respondents in the pandemic survey whether they had children (in the pre-pandemic survey, 20% of postdocs answered that they had children), 10% of respondents mentioned in comments that ensuring their children had proper care was a major stressor and led to severe work disruptions. Additionally, 68% of these comments were from female respondents and 32% from males suggesting a greater burden of childcare for female postdocs. Overall, childcare was the 5th most frequently mentioned stressor (**Figure 2A-B**). Parents mentioned “I have lost childcare for my baby and it has had a significant impact on my ability to write, complete research goals, and apply for grants”; “It was difficult to do any writing- or reading-based work because the daycares were closed, and my partner and I had to divide the day into childcare/work time”; “Loss of productivity due to loss of childcare, feeling like I am slipping behind my colleagues without children”. Some reported feeling burnt out from putting in long hours and mentioned lack of support from their peers and their university; “Lack of childcare and intense pressure from PI to continue long hours at home”; “Loss of childcare and co-workers not respectful of the loss of childcare”; “My institution enacted strict … “shift schedules” that were outside of childcare hours so I was unable to work a full work week. However, I was expected to produce the same (if not more) results/data to make up for the time we were locked out” (more examples in **Table 1**).

Shutdowns also had an adverse impact on postdocs’ relationships with their Principal Investigator (PI) and coworkers. When asked if respondents were able to maintain regular contact with their PI and coworkers, half of the respondents (50%) reported they had but not as much as before the pandemic and 1% reported no contact (49% reported maintaining as much contact as before the pandemic). In open-ended responses, postdocs indicated facing high demands from PIs and unrealistic expectations to be productive during the pandemic (examples in **Table 1**). Some felt that work from home was expected to be “business as usual” and there was immense pressure to “work round the clock”, “work long hours and continuously produce results” and “produce data when no lab activities are allowed”. One respondent indicated inability to utilize institutional support due to over-work: “the PI puts a large amount of pressure and therefore there is really no time to make use of any of the resources”. Conversely, supportive PIs were lauded for their role in lessening stress. Respondents mentioned: “I did not have a lot of stress factors. I was lucky to have a supportive PI that understood how stressful a time this can be and set a pretty low expectation bar”; “working from home during shutdown with a 5yo kid was impossible, really stressful and I am happy my PI was understanding and let me work half time.”

International postdocs reported more difficulty in meeting basic needs such as health care (10% vs. 6%) and food (4% vs. 1%), while US citizens/PR reported more difficulty in obtaining childcare (13% vs 9%) (**Figure 3A**). Additionally, international respondents (n=718) expressed specific worries regarding their residency status. The majority of international postdocs reported apprehension about immigration or visas either due to recent policy changes in the US (84%) or in general (11%) (**Figure 3B**). The primary concerns noted were traveling (75%), US immigration policy changes (69%), and travel bans (68%) (**Figure 3C, Table 1**). Furthermore, more international females than males were worried about immigration issues (89% vs. 78%) (**Figure 3–figure supplement 1A**); specifically, travel (80% vs. 70%), delays in visa renewal (65% vs. 56%), and travel bans (72% vs. 62%) (**Figure 3–figure supplement 1B**).

**Figure 3:**
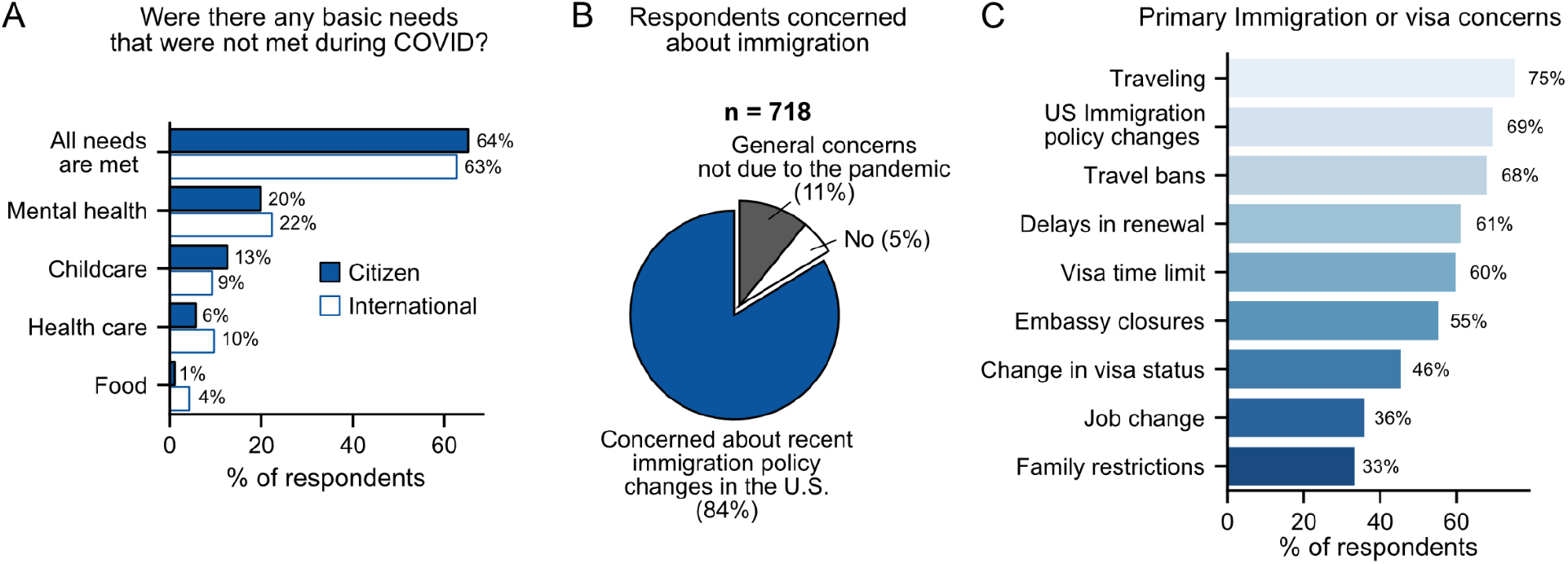
Impact of COVID-19 on international postdocs. **A**. Citizenship status had a significant impact on health care (Chi-squared test, p=0.0023, χ2 = 9.3), childcare (Chi-squared test, p=0.035, χ2 = 4.4) and food (Chi-squared test, p=1.8×10^−5^, χ2 = 9.3) basic needs that were left unmet during the pandemic (n=1,657). **B**. International postdocs’ concerns about immigration and visa (n=718). **C**. Primary immigration or visa concerns (n=718). See **Figure 3–figure supplement 1** for breakdown of immigration concerns by gender.

### Mental Health and Wellness

Overall, 76% of respondents stated that the COVID-19 pandemic had impacted their mental health, with 32% stating that it had a high or very high impact (**Figure 4A**). In open-ended responses, postdocs mentioned significant impacts on their mental health due to isolation and pandemic associated stressors leading to reduced productivity, inability to focus and work effectively: “My mental health has been struggling, which has negative consequences on my ability to focus”; “The isolation has had a negative effect on my mental health …”; “Mental health diminished productivity despite being able to work 100% remotely” (see **Table 1** for more examples).

**Figure 4:**
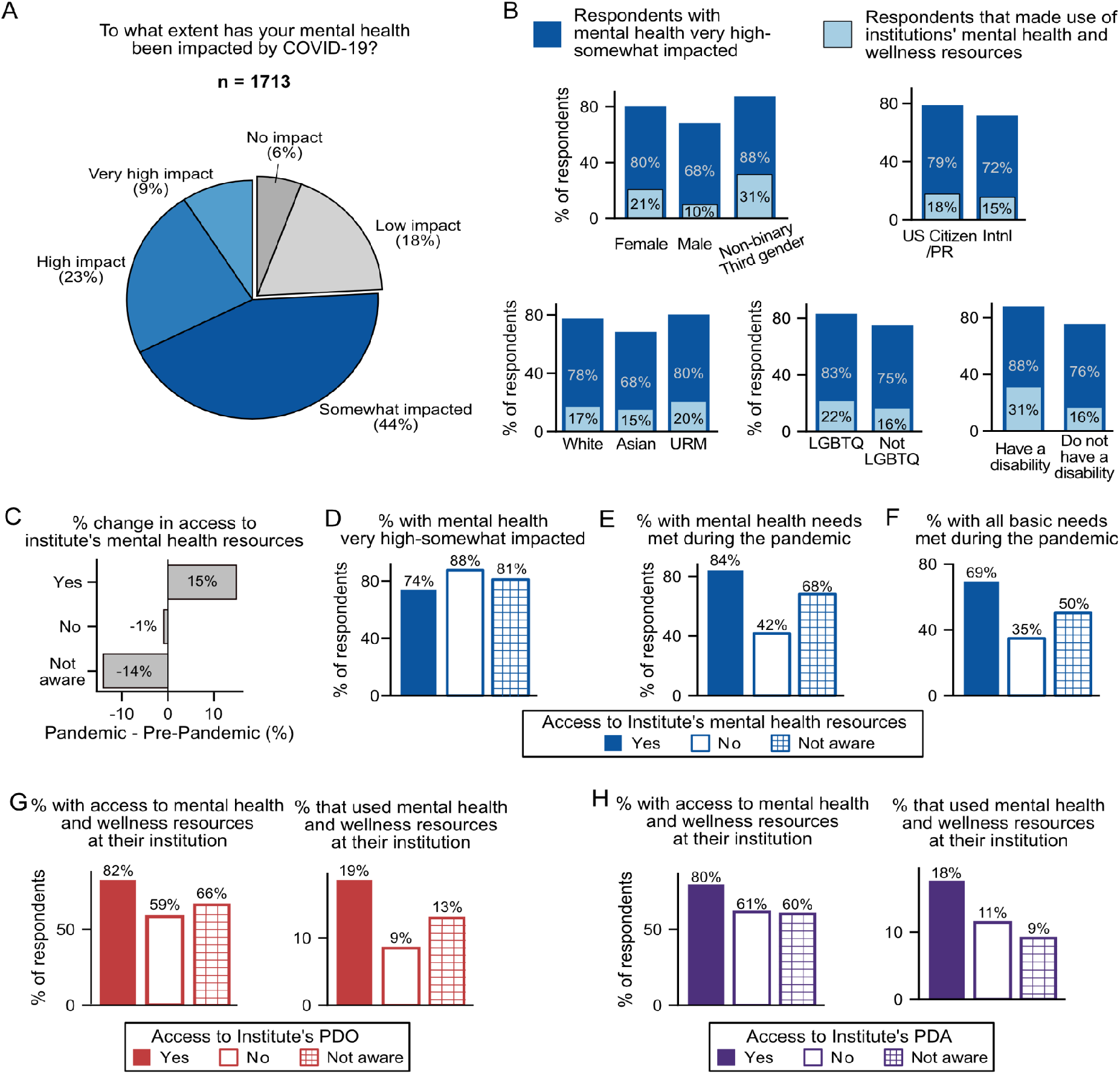
Impact of COVID-19 on mental health. **A**. The majority of survey respondents stated that COVID-19 had impacted their mental health while only 6% stated that it had no impact (n=1,713). **B**. Although most surveyed postdocs stated that their mental health was impacted (very higher impact, high impact, and somewhat impacted), a minority of these postdocs utilized mental health and wellness resources provided by their institution. Females and non-binary/Third gender had more impact than males (n=1,691; ordinal logistic regression OR=0.51,[95% CI:0.43-0.62],p= 1.95e-12 and OR=0.30[95% CI:0.12-0.74],p=0.0085 respectively) and used more institutional resources (Chi-squared test p=2.46×10^−8^, χ2 =35.04), US Citizens/PR reported a greater impact on mental health than International postdocs (n=1,693; ordinal logistic regression, p= 0.0022, OR=1.32[95% CI:1.11-1.58]). Asian postdocs had less impact compared to white (n=1,667; ordinal logistic regression, p= 6.88e-6, OR=0.61,[95% CI:0.49-0.75]) and URM (ordinal logistic regression, p= 1.04×10^−4^, OR=0.54,[95% CI:0.40-0.74]). LGBTQ community (n=1,682; ordinal logistic regression, p= 3.28×10^−5^, OR=2.02,[95% CI:1.45-2.81]) and postdocs with disabilities (n=1,682; ordinal logistic regression, p= 9.66×10^−5^, OR=3.09, [95% CI:1.75-5.44]) reported higher impact on their mental health. Postdocs with disabilities also used more institutional resources (Chi-squared test, p=0.024, χ2 = 5.11) **C**. During the pandemic, more individuals had access to mental health resources, which was reflected in an increased awareness of these resources available at their institution (Chi-squared test, p=3.8×10^−30^, χ2 = 135.5; n pre-pandemic=5,795, n pandemic=1,713). That increase in awareness is proportional to the increase in respondents stating that their institution has available mental health resources. **D**. Having access (ordinal logistic regression, p= 3.54×10^−6^, OR=2.83,[95% CI:1.83-4.40]), or being aware of (ordinal logistic regression, p= 0.011, OR=1.34, [95% CI:1.07-1.67]) mental health resources reduced mental health impact during COVID-19 (n=1,710). **E**. A larger portion of postdocs having access to mental health resources had their mental health basic needs met (Chi-squared test, p=2.18×10^−23^, χ2 = 104.36; n=1,722). **F**. A larger portion of postdocs having access to mental health resources had all their basic needs met (Chi-squared test, p=6.78×10^−16^, χ2 = 69.86; n=1,722). See **Figure 4–figure supplement 1A** for other basic needs unmet vs access to mental health resources. **G and H**. Having access to a PDO or a PDA increased access to (PDO (Chi-squared test, p=6.66×10^−24^, χ2 = 114.87; n=1,697); PDA (Chi-squared test, p=1.39×10^−14^, χ2 =71.01; n=1,703)) and the use (PDO (Chi-squared test, p=0.002, χ2 = 12.32; n=1,694); PDA(Chi-squared test, p=0.016, χ2 = 8.29; n=1,699)) of mental health resources.

All gender, race and ethnicity, and identity groups indicated a significant impact on mental health. However, certain groups reported more of an impact than others; females and third gender/non-binary reported a greater impact than males (80% and 88% vs. 68%); US citizens/PR reported more of an impact than international postdocs (79% vs. 72%); white and URM postdocs reported more of an impact than Asian postdocs (78% and 80% vs. 68%); members of the LGBTQ community (83% vs. 75%) and postdocs with disabilities (88% vs. 76%) reported more of an impact than postdocs not identifying with these groups (**Figure 4B**).

Parallel to this impact on mental health, access to institutional mental health resources rose by 14% (**Figure 4C**), which appears to be linked to an increase in awareness, although only 17% of postdocs indicated use of these resources. Certain groups reported higher usage of these resources: female and third gender/non-binary postdocs compared to male (female 21% and non-binary/third gender 31% vs. male 10%) and postdocs with disabilities compared to those without disabilities (31% vs. 16%, **Figure 4B**). Some of the groups that indicated a greater impact on their mental health (females, third gender/non-binary, postdocs with disabilities) were also more likely to access mental health resources (**Figure 4B**), while other groups that reported a higher impact on mental health (white, LGBTQ and US citizens/PR) were less likely to seek help (**Figure 4B**). Notably, postdocs without access to, or who were unaware of, institutional mental health resources were more likely to have their mental health impacted by COVID-19 than postdocs with those resources (**Figure 4D**). These data suggest: the broad effect of COVID-19 on mental health in US postdocs; indicate unmet needs in this trainee population; and highlight the significance of institutional resources.

Indeed, postdocs were more likely to have their mental health needs met if their institution provided these resources (84%) than if their institution either did not provide them (42%) or if they were unaware of these resources at their institution (68%, **Figure 4E**). Access to institutional mental health resources was also associated with whether postdocs had their basic needs met during the pandemic. Overall, postdocs at institutions that provided mental health resources were more likely to have all their basic needs met (69%) compared to those without (35%) or unaware of these resources (50%) (**Figure 4F**). Unsurprisingly, postdocs that did not have access to, or were unaware of mental health resources at their institutions, were also more likely to have other basic needs unmet such as food (8% (no), 2% (yes), 4% (not aware)) or health care (21% (no), 7% (yes), 7% (not aware); **Figure 4–figure supplement 1A**). In written responses, several postdocs mentioned that their institutions provided mental health resources, however they were often unaffordable or inaccessible: “…doesn’t take postdocs appointments for mental health or other such services they are completely booked [*sic*].”; “Limited financial resources to pay to access mental health resources as “free” sessions through employer was used pre-COVID.”; “ … has mental health resources but they are not free at all.”; “Note, the mental health resources available to post-docs and faculty here are minimal, but they do exist --mostly things like meditation workshops. … However, whether any of these resources are available to postdocs depends on whether our funding is internal (‘associates’, as I am) or external (‘fellows’, who receive fewer benefits)”. These stark differences between institutions with mental health resources and those without, highlight the widespread importance of mental health care and its correlation with quality of life in the postdoctoral population.

As previously indicated (**Figure 2F**), access to a PDA and/or a PDO also increased the likelihood of mental health needs being met. This trend may be due in part to a larger proportion of postdocs with access to a PDO/PDA also having access to mental health resources (82% and 80%) compared to those that did not (59% and 61%) or were unaware (66% and 60%) (**Figure 4G-H**). Postdocs with a PDO/PDA were also more likely to use their institution’s mental health resources (19% and 18%) compared to those that did not have access (9% and 11%) or were unaware of these resources (13% and 9%, **Figure 4G-H**).

### Career Trajectory

The pandemic dramatically impacted career trajectories of the postdocs due to lab shutdowns, inability to communicate with faculty supervisors and research group members, and most significantly, additional family responsibilities, etc., compared to one year earlier (see word cloud in **Figure 2A-B** and select comments in **Table 1**). This resulted in reduced research productivity, delayed job searches, lowered confidence in attaining the desired career, and uncertainty in overall career trajectory. Even though the postdocs were older and had more years of experience when re-surveyed (**Figure 1D-E**), a smaller proportion were currently looking for positions (64% pre-pandemic, 56% during the pandemic), with 11% of postdocs specifically delaying their job search because of the pandemic (**Figure 5A**). In addition, postdocs were less confident in achieving their career goals than before the pandemic (**Figure 5B**), which may be contributing to the observed decline in those actively pursuing new positions (**Figure 5A**). Furthermore, more postdocs were undecided about their future careers than before the pandemic (9% to 12%) (**Figure 5C**). Together, these results highlight the substantial increase in career uncertainty felt by postdocs.

**Figure 5:**
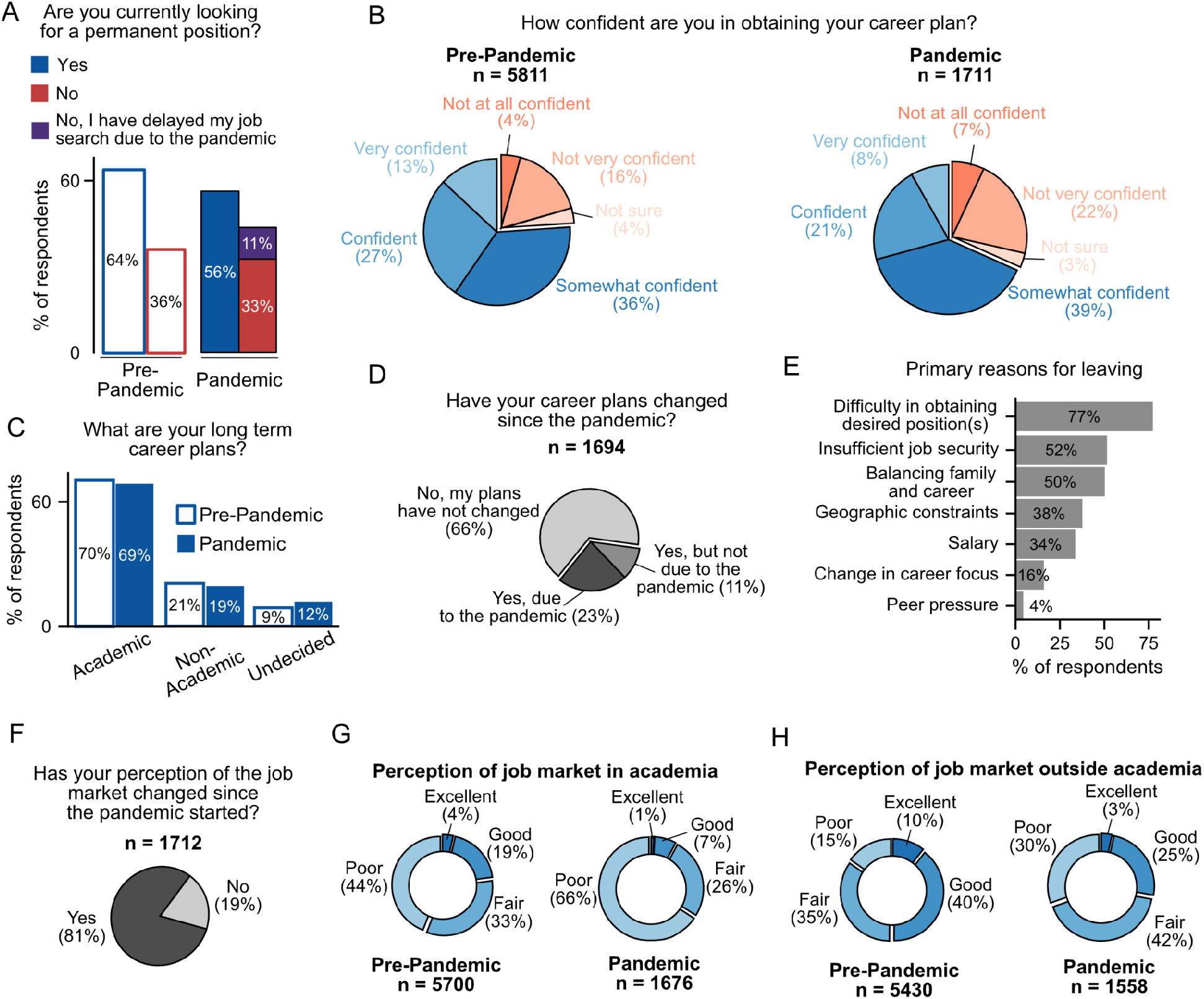
The effect of COVID-19 on career trajectories of postdocs. **A**. Fewer postdocs are actively looking for a permanent position (n=1,704) than before the pandemic (n=5,676; Chi-squared test, p=2.1×10^−8^, χ2 = 31.39). See **Figure 5–figure supplement 1A** for breakdown by type of position. **B**. Postdocs are less confident in their ability to obtain their desired career since the start of the pandemic (ordinal logistic regression, p= 1.58×10^−20^, OR=0.62,[95% CI:0.56-0.69]; n pre-pandemic=5,811, n pandemic= 1,711). **C**. The long-term goals of postdocs have not shifted during the pandemic. However, a larger proportion of postdocs are now uncertain about their career trajectories (Chi-squared test, p=0.0022, χ2 = 41.3; n pre-pandemic=5,746, n pandemic= 1,716). **D**. 34% of postdocs indicated that their career plans changed since the pandemic started (n=1,694). **E**. Primary reasons for changes in career trajectory (n=388). See **Figure 5–figure supplement 1B-C** for breakdown by citizenship status and race/ethnicity. **F**. During the pandemic, the perception of both the academic and non-academic job markets has declined (n=1,712). See **Figure 5–figure supplement 1D-E** for breakdown by citizenship status and race/ethnicity. **G**. A decrease in the perception of the job market both in (ordinal logistic regression, p=2.32×10^−63^, OR=0.39,[95% CI:0.35-0.43]; pre-pandemic=5,700, n pandemic= 1,676)) and **H**. outside (ordinal logistic regression, p=6.5×10^−70^, OR=0.39, [95% CI:0.35-0.43]; pre-pandemic=5,430, n pandemic= 1,558)) academia was observed during the pandemic compared to the pre-pandemic survey.

Overall, 34% of postdocs reported changing their career plans during the pandemic, with 23% of respondents indicating that COVID-19 was the direct cause of their change (**Figure 5D**). This latter group was more likely to be undecided about future careers (20% vs. 7%) or considering non-academic positions (28% vs. 14%), and much less likely to be seeking an academic position (51% vs. 79%) compared to postdocs who did not change their career plans (66% of surveyed postdocs) (**Figure 5–figure supplement 1A**). The main reasons cited for career trajectory changes were: i) difficulty in obtaining the desired position (77%), ii) insufficient job security (52%), and iii) balancing family and career (50%) (**Figure 5E**). Additionally, reasons for career change differed by citizenship status and race/ethnicity. International postdocs cited more peer pressure than US citizen/PR (8% vs. 1%), while the latter noted more difficulty in obtaining desired positions (83% vs. 69%) as well as balancing family and career (58% vs. 39%, **Figure 5– figure supplement 1B**). Moreover, Asian postdocs indicated more peer pressure as a reason for changing career trajectory (9% vs. 4% in URM and 3% in white, **Figure 5– figure supplement 1C**). Lastly, we observed no differences by gender or identity groups with respect to reasons for changing career trajectory (Chi-squared test, p>0.05, data not shown).

The majority of postdocs surveyed also reported a change in their perception of the job market (81%) (**Figure 5F**), with certain subgroups reporting differential changes; more US citizens/PR than international postdocs (85% vs. 74%, **Figure 5–figure supplement 1D**) and fewer Asian (77% compared to URM (83%) and white (82%), **Figure 5–figure supplement 1E**) reported a change in perception. No differences were observed based on gender or identity groups (Chi-squared test, p>0.05, data not shown). This altered perception was observed for both the academic and non-academic job markets. Overall, the majority of the respondents viewed the current academic job market as poor (66%) or fair (26%), which is a significant change compared to the pre-pandemic survey, where fewer postdocs viewed the market as poor (44%) and more viewed it as fair (33%). Although the perception of the job market outside of academia was better −28% of the respondents found it either excellent or good compared to academic careers (8%) - there was still a decrease in perception from the pre-pandemic survey (**Figure 5G-H**). Altogether, the perception of both career paths had markedly declined (**Figure 5G-H**).

### Career Changes During the Pandemic

The postdoctoral position is considered temporary with the ultimate goal of providing the necessary training and experience to successfully transition to more permanent careers. To better understand the effects of the pandemic on career outcomes, we surveyed those who were no longer in postdoctoral positions. Of those who responded to the second survey, 11% (219/1,941) were no longer postdocs, with 14% indicating that this career transition was a consequence of the pandemic (**Figure 6A**). Overall, 56% of the postdocs who made career transitions remained in academic positions (clinical, research staff, or faculty), while nearly 8% were unemployed. When we separately examined the postdocs who made career transitions as a consequence or irrespective of the pandemic, we observed a profound difference in career outcomes. The former group was more likely to be unemployed (38% vs. 6%) and less likely to be in academic positions than postdocs who chose to leave their position regardless of the pandemic (24% vs. 65%), while we observed little difference in those pursuing non-academic careers (38% vs. 29%; **Figure 6B**).

**Figure 6:**
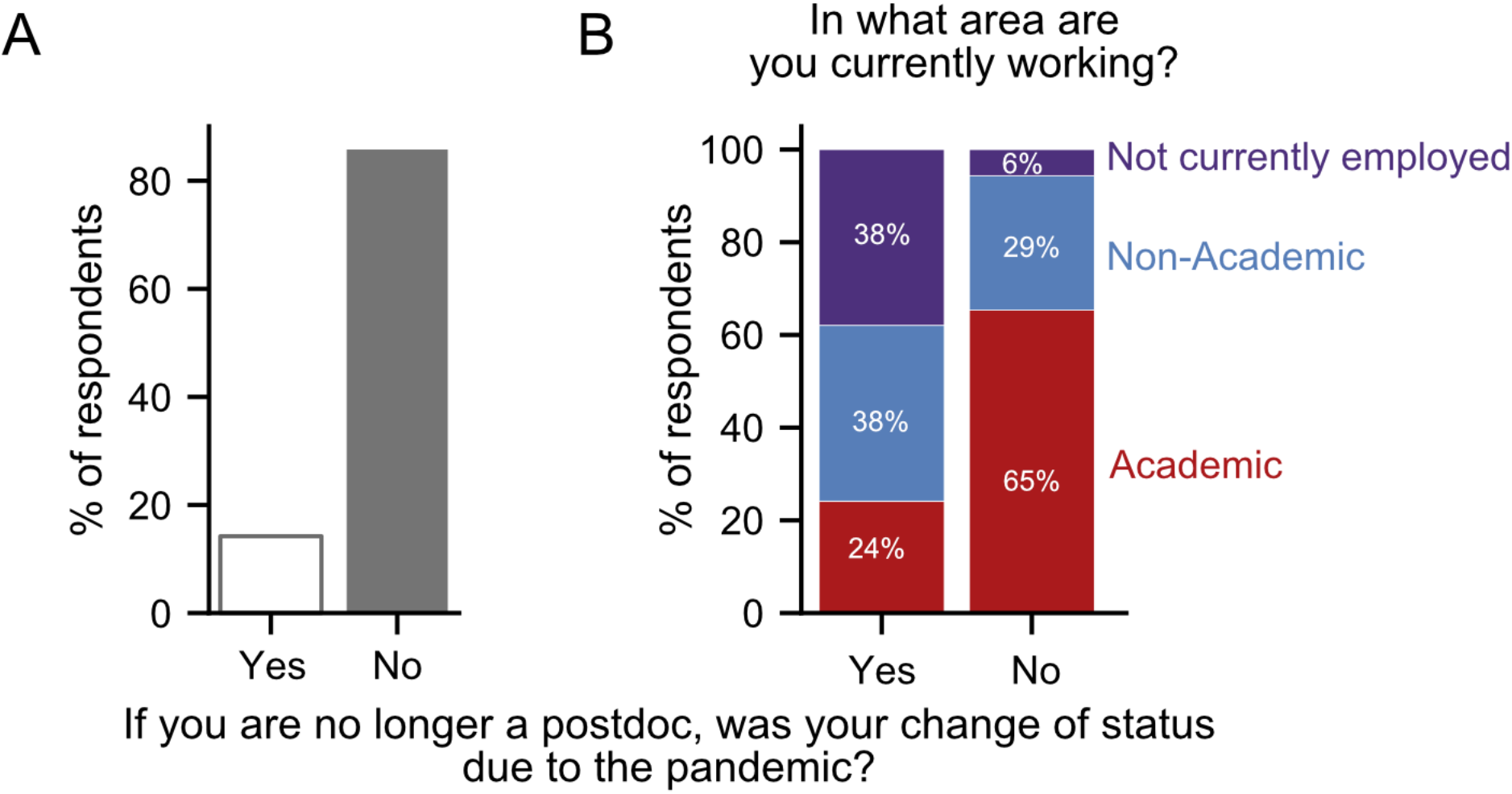
Career transitions made during the pandemic. **A**. 14% of respondents who indicated that they are no longer a postdoc, stated that their transition was a consequence of the pandemic (n=218). **B**. Postdocs who transitioned due to the pandemic were more likely to be unemployed (purple) and less likely to have an academic position (red) than postdocs whose transition was not a consequence of the pandemic (Chi-squared test, p=6.69×10^−8^, χ2 = 33.04; n=205).

## Discussion

Early in March of 2020, the COVID-19 pandemic forced research facilities across the US to drastically alter their activities. This resulted in a cascade of events, including loss of research progress, career advancement, and a further imbalance of work and life activities. To investigate the impact of these changes on the postdoctoral experience, we took advantage of our recently completed national postdoctoral survey (June - December 2019) and re-surveyed the same population during the pandemic (between October 1 and November 3 2020). Unsurprisingly, given that the pandemic survey was conducted in a subset of the pre-pandemic survey, the demographics were comparable between the two surveys, with the exception of the respondents being older and further along in their careers, as expected. Furthermore, as the survey was only open during a restricted period (1 month), it allowed us to capture a defined period of the pandemic. Even though we did not interrogate during the first few months with full lockdowns, we surveyed postdoc during the second wave (in the US), when many institutions were only partially opened to support social distancing, before access to vaccines and right before the 2020 US elections. Our data provide a unique opportunity to directly assess the effects of the pandemic on the postdoctoral experience.

Although there have been multiple reports of the pandemic’s impact on the STEM workforce^13,22–25^ few have discussed postdocs specifically^26,27^. Using our pandemic survey, we were able to ascertain the impact of COVID-19 on mental health, ability to meet basic needs, and career trajectory; as well, the analysis revealed the importance of institutional resources for postdocs. Although our surveys indicate that the majority of all postdocs were affected by loss of productivity and overwhelming mental health challenges during the pandemic, demographic subgroups experienced the effects of the pandemic differently. Furthermore, our survey highlights the additional burden of the pandemic on international postdocs, those from underrepresented minority groups, and women. Importantly, the ability to do a comparative analysis of pre-pandemic to pandemic responses revealed profound effects of the pandemic on career trajectories of postdocs. Many postdocs also provided commentary to the two open-ended response questions in our pandemic survey (2,768 comments were collected), which further demonstrated the impact of the pandemic on the postdoctoral population (see representative quotes in **Table 1**).

As previously indicated, this survey provides a unique “before-and-during” opportunity to observe the effects of COVID-19 on postdoctoral life. However, there were some limitations to our study. First, although the pandemic survey was conducted in a subset of the pre-pandemic respondents and therefore was more directly comparable, the responses were anonymized, and we are unable to do a direct one-to-one comparison of pre-pandemic to pandemic responses on an individual level. Furthermore, although we were able to assess caregivers through responses to a handful of questions, including the written responses, we did not directly ask if respondents were parents or caregivers, limiting our ability to assess those effects more directly. Lastly, because of sample sizes, we were limited in our ability to evaluate certain metrics for some demographics such as the LGBTQ, third gender, individuals with disabilities and certain races/ethnicities. To be able to parse out potential differences between racial and ethnic groups, we pooled all individuals into three broad groups; white, Asian and URM. Nonetheless, these data still represent a rich collection of information about the postdoctoral experience before and during the pandemic.

As is apparent from our survey data, access to institutional resources is critical not only for the ability of postdocs to complete their work in safe and supportive environments - as is often the focus of institutional efforts - but also for their mental and physical wellbeing as we note in this manuscript. Along with these resources, our data indicate the importance of institutional tracking of postdoc populations. As we previously reported^20^, postdocs are an often overlooked and forgotten population in academia, with a non-negligible number of institutes being unaware of their total postdoc population, let alone the concerns of that population. Here we’ve shown that nearly a quarter of all postdocs felt that their mental health needs were unmet during the pandemic and just as unsettling, a non-negligible proportion struggled with access to food (2%) and healthcare (7%). In a position that emphasizes sacrifice for research, institutions need to pay more attention to ensure that minimal basic needs are met and assume responsibility for these burdens. Moreover, respondents that were no longer in postdoctoral positions due to the pandemic had higher rates of unemployment. We did not collect detailed information about these former postdocs and more follow-up studies are needed to track their outcomes. Furthermore, in the ~7 months between the beginning of the pandemic and the survey, we were already able to see hints of long-term consequences such as delayed job searches, lost productivity, lost positions, fewer opportunities, and altered career trajectories. Moving forward, we plan to continue to survey US-based postdocs in order to generate a better understanding of the long-term consequences of the COVID-19 pandemic on postdoc experiences and outcomes. Ultimately, understanding the needs of this critical workforce will also broadly benefit the future of science and research.

## Methods

### Survey design and dissemination

The National Postdoctoral Survey was designed to capture the experiences and demographic information of postdoctoral fellows and scholars across the United States. The survey was initially conceived and developed by postdocs within the University of Chicago’s Biological Sciences Division Postdoctoral Association (PDA) in 2016, in order to identify important issues within the postdoctoral community and inform and equip those who advocate for postdoctoral policies to make positive changes. The results of the first National Postdoc Survey were published by McConnell, et al. in 2018^20^.

In 2019, a second updated version of the National Postdoc Survey was launched by the University of Chicago PDA. This version, referred to as the “pre-pandemic survey”, collected responses from postdocs in the United States from June 4, 2019, until December 31, 2019. In order to make postdocs across the US aware of the survey, multiple types of grass-roots outreach were used in a similar manner to McConnell, et al.^20^ First, we performed online website searches for Postdoc Offices (PDOs) at doctoral degree-granting universities or research institutions in the US that train postdocs. We compiled a list of publicly available email addresses for institutional representatives of these PDOs. If we were unable to identify a PDO, or if an institution did not have a PDO, then contact information was collected instead for an administrative or faculty representative within an Office of Research, Graduate School, or Provost, or for a similar official who might have access to postdocs. We also collected contact information, if available, for postdoc leaders of Postdoctoral Associations (PDA), which we contacted if institutions did respond to our initial outreach or if an institution’s response rate was deemed low compared to the 2016 National Postdoc Survey. We emailed over 400 institutional PDOs, other administrative contacts or PDA leaders, described the goals of the survey, and asked them to distribute our survey link and invitation to the postdocs at their institution. Over the course of the 7 months that the survey was open, follow-up emails were sent to our contacts to remind them to send the email to their postdocs, or to distribute the survey link if they had not already done so. In addition to our outreach to institutional representatives, we shared the survey on social media websites including Twitter and LinkedIn, launched a website dedicated to the National Postdoc Survey, and prepared an email campaign to advertise the survey which was distributed by the National Postdoctoral Association to its large national listserv of postdocs and postdoc advocates. These additional methods were used to enhance awareness of the survey and distribute the survey link directly to postdocs who may not have received it through their institution.

During the 7 months that the survey was open, responses from 6,292 postdocs were collected from over 300 institutions in nearly every state in the nation. All responses were collected anonymously, but many respondents voluntarily provided contact information in a separate form to draw names for survey incentive prizes. Of the 6,292 respondents to the survey, 5,929 identified as postdocs at a US institution and only their responses were used for analysis.

While analysis of the 2019 pre-pandemic survey data was underway, the COVID-19 pandemic commenced, and it became evident that a follow-up survey was necessary to assess the changes brought on by the pandemic in the mindsets and current situations of postdocs. Questions were designed in 2020 for a shorter “pandemic survey” to query what changes the postdocs experienced in their career goals and whether their plans changed since the pandemic started, current perceptions of the job market in academia, and how their research and life has been affected by the pandemic. All postdocs who completed the initial pre-pandemic survey and submitted their email addresses for recontact were asked to complete this second pandemic survey, which was launched on October 1, 2020 and stayed open for one month. In total, 1,942 responses to the pandemic survey were collected. Of these responses, 1,722 were submitted by researchers currently in postdoctoral positions in the United States, and these responses are analyzed here. Pre-pandemic and pandemic survey questionnaires are included in **supplementary file 1** and **supplementary file 2** respectively.

### Data Analyses

We used two definitions of race and ethnicity, a more granular one: comparing each group to the rest of the respondents (white/Caucasian, Asian/Asian American, South Asian/South East Asian, Black/African American, Hispanics/Latinos, Middle Eastern, Native American/Alaska Native, and Pacific Islander/Hawaiian Native), and a more consolidated one classifying samples in three groups (underrepresented minority (URM): Black/African American, Hispanics/Latinos, Native American/Alaska Native, and Pacific Islander/Hawaii Native); Asians (Asian/Asian American and South Asian/SouthEast Asian); and white (white/Caucasian and Middle Eastern)).

Non-respondents were removed before each analysis. To assess differences, we used either ordinal logistic regression in the presence of ordinal dependent variables (using the R package “MASS”) or Chi-square test in the presence of categorical data (basic R function). We considered p-values <0.05 to be significant. In the manuscript, p-values of <0.05 were identified as *, p<0.01 ** and p<0.001 ***. Word clouds were generated in Python using the wordcloud package. Figures were generated using Python version 3.7.6.

## Acknowledgements

We thank Dr. Valerie Miller (UIC) for her assistance with the 2019 survey instrument design and dissemination; the Future of Research directors for suggesting the inclusion of mental health and wellness queries in the survey; and we are especially grateful to Drs. Erin Heckler (Yale University) and Imogen Hurley (UW Madison) for valuable comments. We are very grateful to the many Postdoctoral Associations, administrators, faculty directors, and others who helped distribute our survey to postdocs across the country. Finally, we wish to especially express our gratitude to the postdocs who shared their experiences with us by participating in the surveys.

## Ethics

Human subjects: Participation in this survey was completely voluntary. In the introduction to this survey, we informed the participants of its purpose, and that results of the survey would be disseminated, in aggregate. All responses were recorded in a secure RedCap Database, so they could not be traced back to individual respondents. Responses were combined for data analysis to maintain respondent anonymity throughout data analysis. Our survey design and dissemination protocol was approved by the University of Chicago Institutional Review Board, IRB Protocol Number 15-1724.

## Data Availability

Non-privileged data used in this study are available in supplemental tables and additional material related to this manuscript. Due to their sensitive nature, much of the raw data is privileged to prevent individual identification in accordance with IRB protocol. However, summary data for institutions, fields, and regions with more than 50 respondents are available upon request.

## Author details

Andréanne Morin is in the Department of Human Genetics, University of Chicago, Chicago, IL.

Contribution: Conceptualization, Methodology, Formal analysis, Investigation, Writing – original draft preparation, Writing – review & editing, Visualization

For correspondence

Competing interests: None

Britney A. Helling is in the Department of Human Genetics, University of Chicago, Chicago, IL.

Contribution: Conceptualization, Methodology, Formal analysis, Writing – original draft preparation; Writing – review & editing

Competing interests: None

Seetha Krishnan is in the Department of Neurobiology and Institute for Neuroscience, University of Chicago, Chicago, IL.

Contribution: Conceptualization, Methodology, Formal analysis, Investigation, Visualization, Writing – original draft preparation, Writing - review & editing

Competing interests: None

Laurie E. Risner is in the Department of Pediatrics, University of Chicago, Chicago, IL.

Contribution: Conceptualization; Methodology; Writing – original draft preparation; Writing – review & editing; Project administration

Competing interests: None.

Nykia D. Walker is in the Ben May Department for Cancer Research, University of Chicago, Chicago, IL.; current affiliation: the University of Maryland, Baltimore County

Contribution: Writing-review & editing

Competing interests: None

Nancy B. Schwartz is in the Department of Pediatrics and the Department of Biochemistry and Molecular Biology, University of Chicago, Chicago, IL.

Contribution: Conceptualization; Methodology; Investigation; Writing – original draft preparation; Writing – review & editing; Visualization; Supervision; Project administration

Competing interests: None

For correspondence

## Supplementary Information

**1. Supplementary file 1: Pre-pandemic survey questionnaire**

**2. Supplementary file 2: Pandemic survey questionnaire**

**Supplementary file 3:** Race and ethnicity distribution among respondents of the prepandemic and pandemic survey.

| **Pandemic Survey** |  |  |  |  |  |  |  |  |
| --- | --- | --- | --- | --- | --- | --- | --- | --- |
|  | Hispanic/<br>Latino | White/<br>Caucasia<br>n | Black/<br>African<br>America<br>n | Asian /<br>Asian<br>America<br>n | South<br>Asian/<br>South<br>East<br>Asian | Middle<br>Eastern | Native<br>American<br>/ Alaska<br>Native | Pacific<br>Islander/<br>Hawaii<br>Native |
| Hispanic/<br>Latino | 89<br>(5.17%) | 62<br>(3.6%) | 11<br>(0.64%) | 2<br>(0.12%) | 0 (0%) | 0 (0%) | 3 (0.17%) | 0 (0%) |
| White/<br>Caucasia<br>n |  | 1016<br>(59%) | 5<br>(0.29%) | 19<br>(1.1%) | 2<br>(0.12%) | 21<br>(1.22%) | 11<br>(0.64%) | 0 (0%) |
| Black/<br>African<br>America<br>n |  |  | 31<br>(1.8%) | 0 (0%) | 0 (0%) | 0 (0%) | 0 (0%) | 0 (0%) |
| Asian /<br>Asian<br>America<br>n |  |  |  | 281<br>(16.32%) | 11<br>(0.64%) | 2<br>(0.12%) | 0 (0%) | 3<br>(0.17%) |
| South<br>Asian/<br>South<br>East<br>Asian |  |  |  |  | 69<br>(4.01%) | 0 (0%) | 0 (0%) | 0 (0%) |
| Middle<br>Eastern |  |  |  |  |  | 27<br>(1.57%) | 0 (0%) | 0 (0%) |
| Native<br>America<br>n/ Alaska<br>Native |  |  |  |  |  |  | 0 (0%) | 0 (0%) |
| Pacific<br>Islander/<br>Hawaii<br>Native |  |  |  |  |  |  |  | 0 (0%) |
| Total* | 174 | 1144 | 51 | 319 | 82 | 52 | 17 | 4 |
| **Pre-pandemic survey** |  |  |  |  |  |  |  |  |

|  | Hispanic/<br>Latino | White/<br>Caucasian | Black/<br>African<br>American | Asian /<br>Asian<br>American | South<br>Asian/<br>South<br>East<br>Asian | Middle<br>Eastern | Native<br>American<br>/ Alaska<br>Native | Pacific<br>Islander/<br>Hawaii<br>Native |
| --- | --- | --- | --- | --- | --- | --- | --- | --- |
| Hispanic/<br>Latino | 312<br>(5.11%) | 188<br>(3.08%) | 13<br>(0.21%) | 4<br>(0.07%) | 0 (0%) | 0 (0%) | 3 (0.05%) | 0 (0%) |
| White/<br>Caucasian |  | 3190<br>(52.20%) | 19<br>(0.3%) | 55<br>(0.90%) | 7<br>(0.11%) | 74<br>(1.21%) | 18<br>(0.29%) | 1<br>(0.02%) |
| Black/<br>African<br>American |  |  | 131<br>(2.14%) | 1<br>(0.02%) | 0 (0%) | 1<br>(0.02%) | 0 (0%) | 0 (0%) |
| Asian /<br>Asian<br>American |  |  |  | 1151<br>(18.83%) | 29<br>(0.47%) | 0 (0%) | 0 (0%) | 9<br>(0.15%) |
| South<br>Asian/<br>South<br>East<br>Asian |  |  |  |  | 291<br>(4.76%) | 0 (0%) | 0 (0%) | 1<br>(0.02%) |
| Middle<br>Eastern |  |  |  |  |  | 124<br>(2.03%) | 0 (0%) | 0 (0%) |
| Native<br>American<br>/ Alaska<br>Native |  |  |  |  |  |  | 2 (0.03%) | 0 (0%) |
| Pacific<br>Islander/<br>Hawaii<br>Native |  |  |  |  |  |  |  | 1<br>(0.02%) |
| Total* | 535 | 3570 | 176 | 1258 | 332 | 206 | 30 | 14 |

In the pandemic survey, 49 (2.85%) respondents did not identify any race or ethnicity and 8 (0.46%) respondents identified $\geq 3$ races or ethnicities. In the pre-pandemic survey, considering only US Postdocs, 301 (4.8%) respondents did not identify any race or ethnicity and 25 (0.4%) respondents identified $\geq 3$ races or ethnicities. \* Number of individual who identify to each ethnicity/race.

**Figure 1–figure supplement 1.**
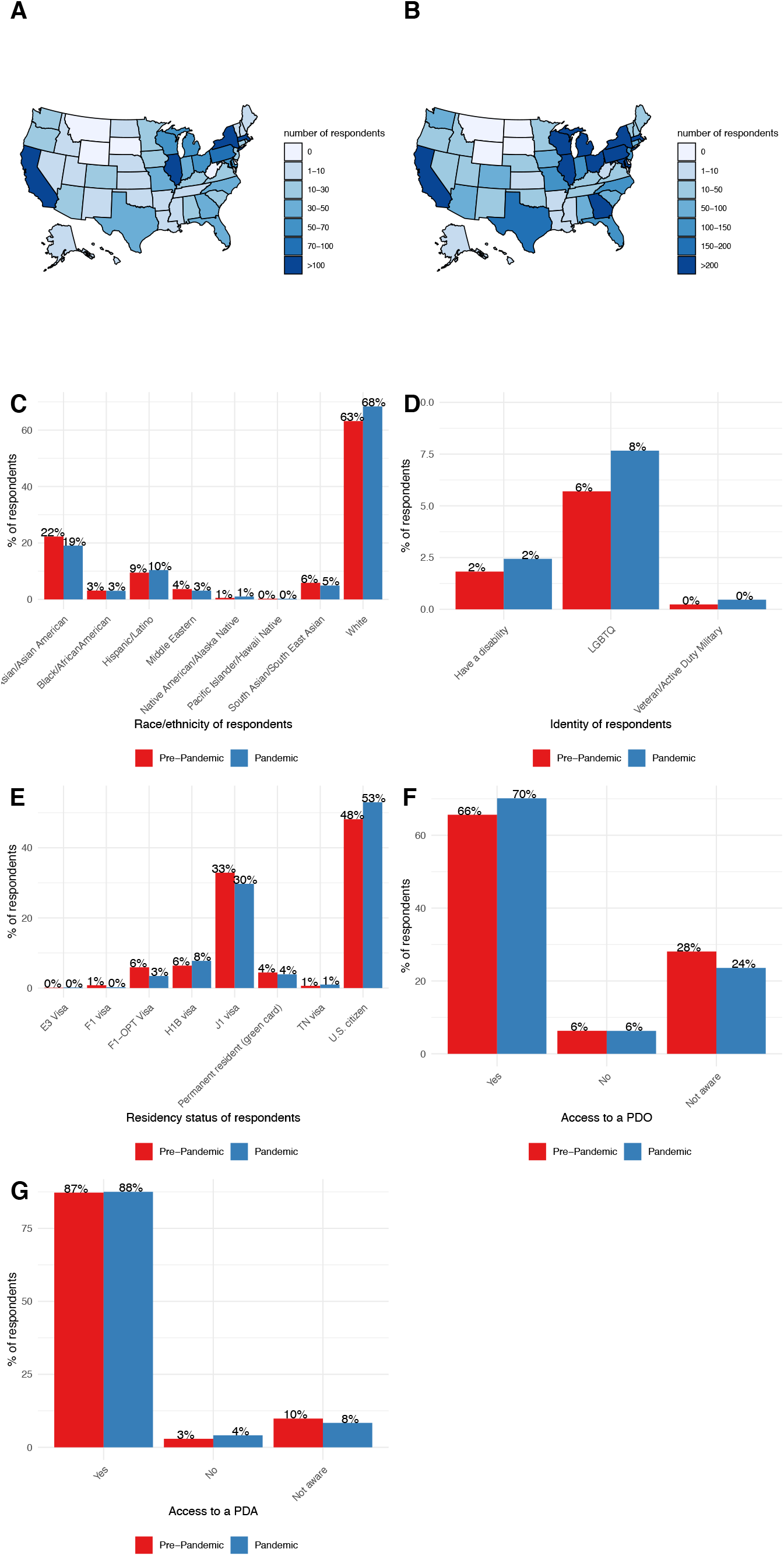
Comparison of demographics between pandemic and prepandemic surveys. **(A-B)** Number of respondents in the pandemic **(A)** and pre-pandemic survey **(B)** by states. **C**. Percentage of respondents by race and ethnicity groups in the pandemic and pre-pandemic surveys. Less respondents identify as Asian and Asian American in the pandemic survey (Chi-squared test, χ2=20.11, p=0.0053). **D**. Percentage of respondents by identity. All of the identity groups were more represented in the pandemic survey compared to the pre-pandemic survey. **E**. Percentage of respondents by residency status, a larger percentage of respondents were US citizens and a smaller percentage of F1-OPT visa holders in the pandemic survey (Chi-squared test, χ2=36.94, p = 1.18×10^−5^). **F**. Increased access to a PDO was observed during the pandemic, mainly due to an increase of awareness of such institutional resource (Chi-squared test, χ2=13.87, p = 9.73×10^−4^). **G**. No differences were observed in access to a PDA before or during the pandemic.

**Figure 2–figure supplement 1.**
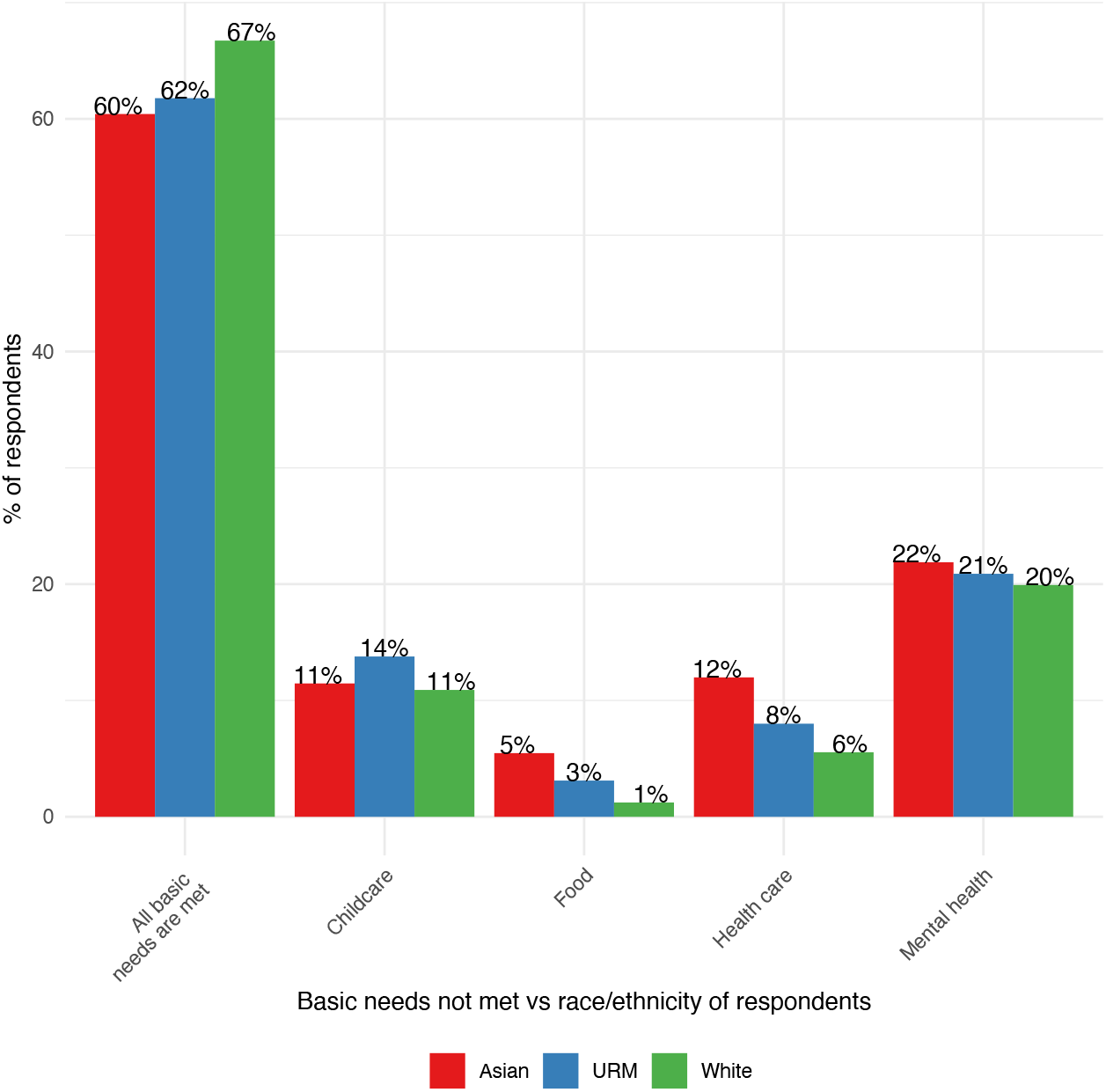
Basic needs not met by race/ethnicity groups. **A**. Postdocs who identified as Asian did not have health care (12% vs 5%, Chi-squared test, χ2=17.3, p=1.7×10^−4^) or food (5% vs 1%, Chi-squared test, χ2=21.76, p=1.88×10^−5^) basic needs met compared to white postdocs.

**Figure 3–figure supplement 1.**
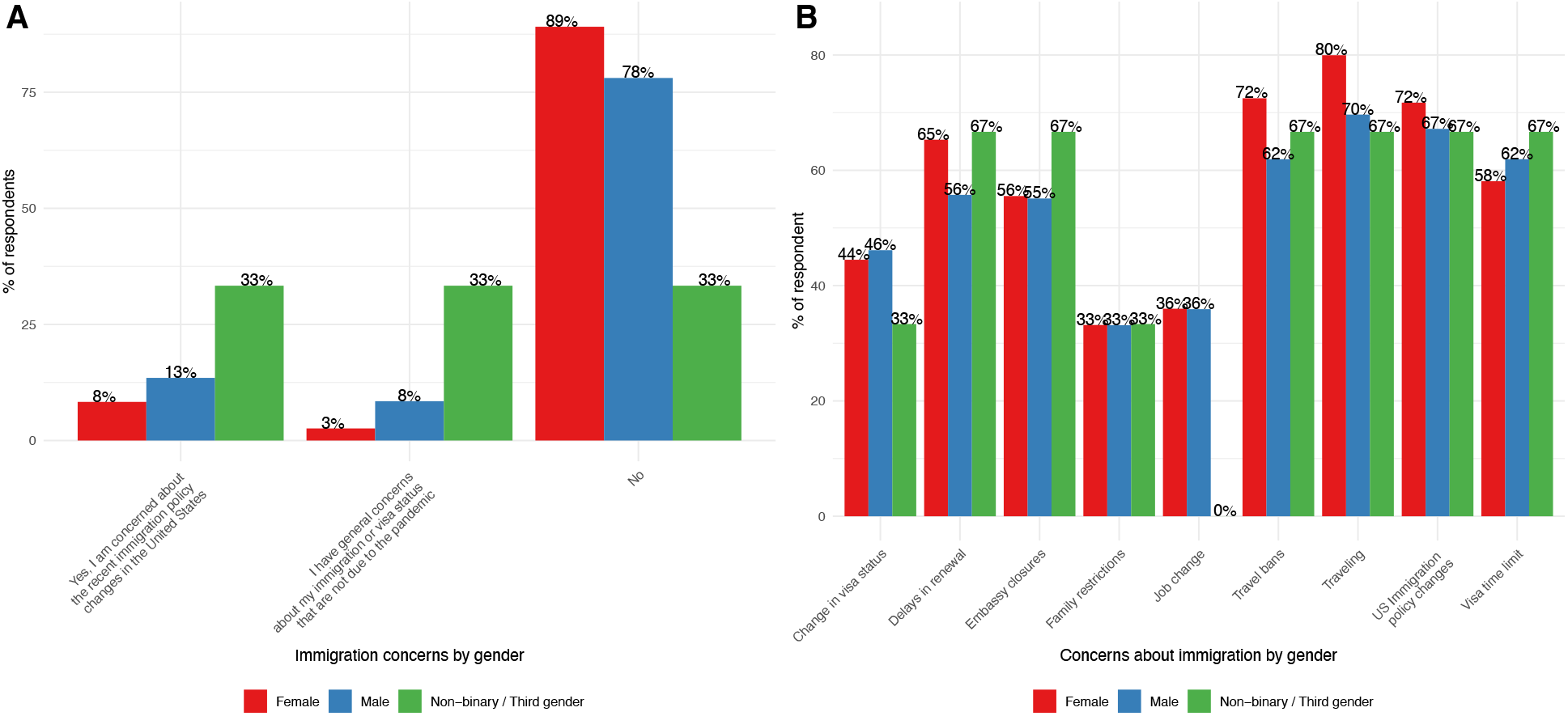
Immigration concerns by gender. **A**. Females were more concerned than males (Chi-squared test, χ2=24.8, p=5.6×10-5) (n=718) and **B**. were more concerned about traveling (Chi-squared test, χ2=10.15, p=0.006), delays in visa renewal (Chi-squared test, χ2=6.83, p=0.032) and travel bans (Chi-squared test, χ2=9.02, p=0.011) (n=715).

**Figure 4–figure supplement 1.**
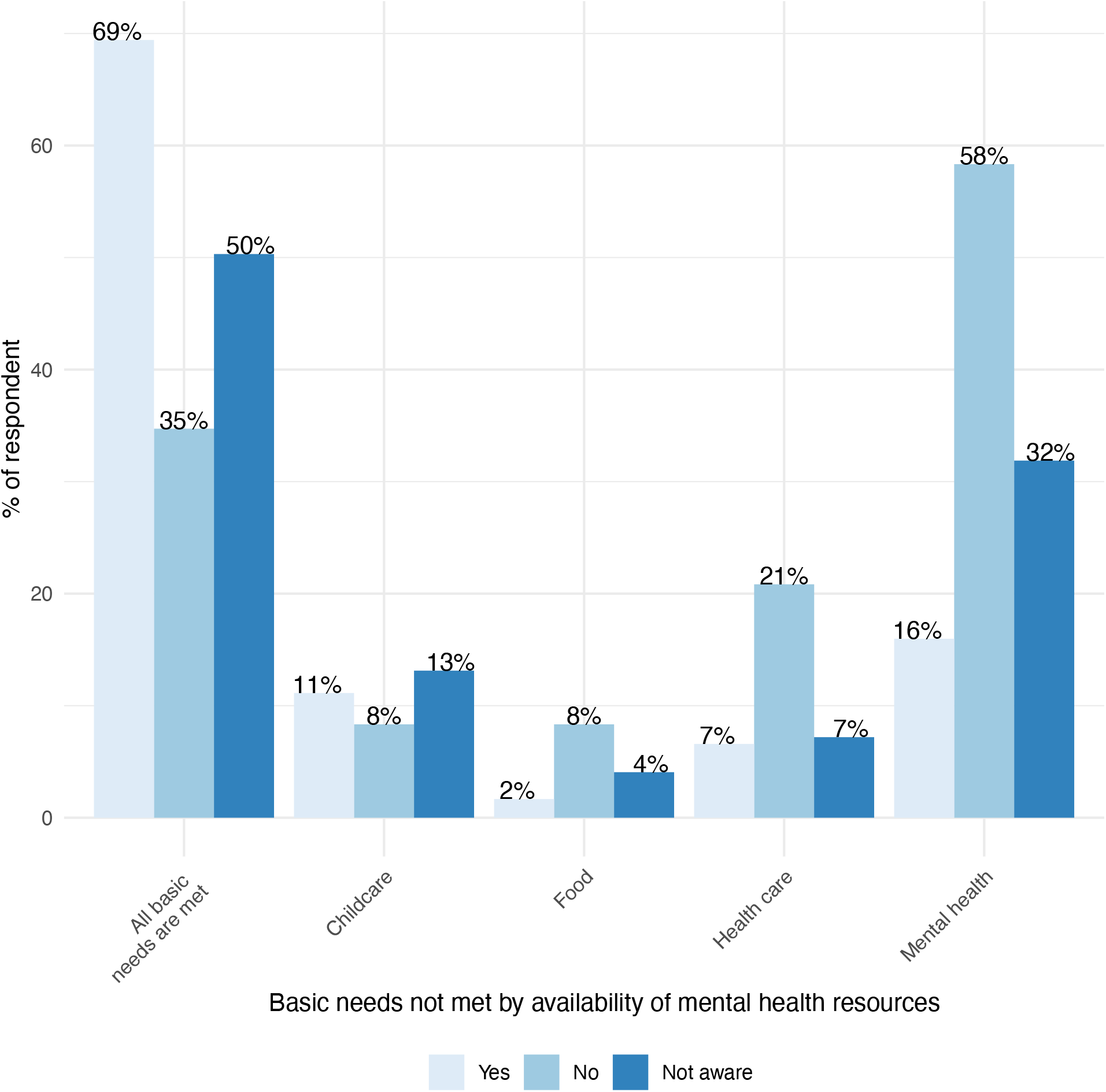
Effect of institutional resources on having mental health needs met. **A**. Postdocs that did not have access to mental health resources through their institutions or were unaware if their institutions had mental health resources were also more likely to have other basic needs unmet such as food (Chi-squared test, χ2=20.5, p=3.54e-5) or health care (Chi-squared test, χ2=17.7, p=1.44e-4).

**Figure 5–figure supplement 1.**
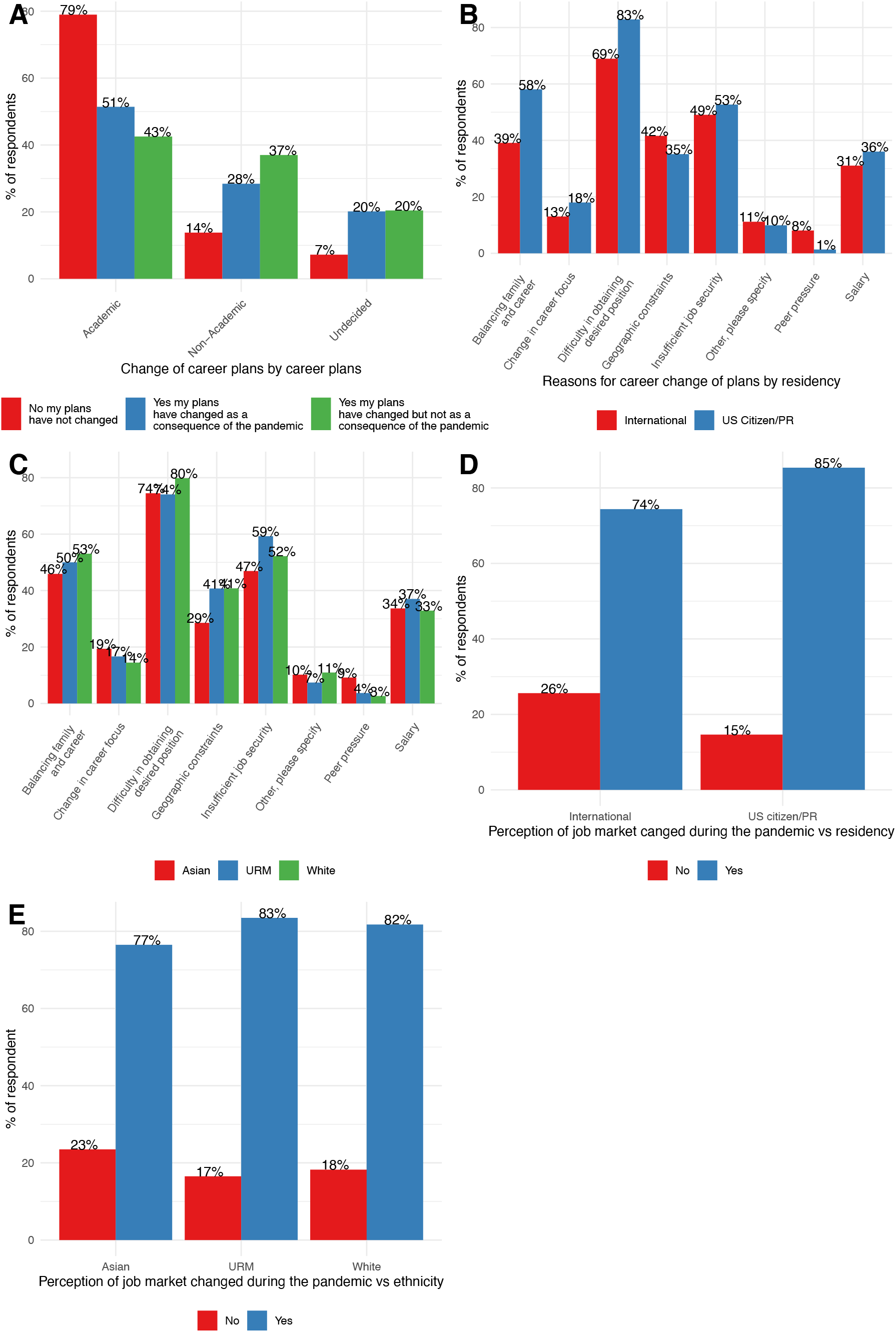
Change in career plans broken down by demographics. **A**. Postdocs that changed their career plans due to the pandemic or not, were less likely to pursue an academic position and were more likely to be undecided (Chi-squared test, χ2=169.91, p=1.09e-35; n=1691). **B**. Reasons for change of career plans differ by residency status (Chi-squared test, χ2=8.92,p=0.0028 (peer pressure), χ2=9.47, p=0.002 (difficulty of finding desired position), χ2=12.7, p=0.00037 (balancing family and career);n=383) and **C**. race/ethnicity (Chi-squared test, χ2=6.97, p=0.031, posthoc p=0.05 (peer pressure);n=380) **D**. Job market perception changed during COVID-19 by residency status, (Chi-squared test, χ2=31.32, p=2.18e-8; n=1,692) and **E**. race/ethnicity, (Chi-squared test, χ2=6.22, p=0.045; n=1,665).

